# Why PAR is not RNA: ion atmospheres, bridging interactions, and ion-induced structural collapse

**DOI:** 10.1101/2025.11.29.691228

**Authors:** Lipika Baidya, Heyang Zhang, Hung T Nguyen

## Abstract

Poly(ADP-ribose) (PAR) is a highly charged, intrinsically flexible nucleic-acid-like homopolymer composed of ADP-ribose units. It serves as a critical post-translational modification that rapidly synthesized in response to DNA damage, and participates in diverse important cellular processes, including DNA repair, chromatin remodeling, and RNA biogenesis, often by promoting biomolecular phase separation. However, PAR’s conformational heterogeneity coupled with limited experimental characterization has hindered a mechanistic understanding of these processes. To address this challenge, we develop a five-bead coarse-grained model of PAR and perform molecular dynamics simulations to compare its conformational ensemble and ion atmosphere with those of RNA across diverse ion environments. Our simulations reveal that PAR undergoes a markedly stronger structural transition than RNA in response to divalent ions, driven by its preferential ion binding and effective ion-mediated bridging between phosphate groups. These interactions facilitate cooperative conformational changes and produce a compact, ion-rich atmosphere that is fundamentally distinct from RNA. Together, our work provides a physically grounded model for PAR, uncovers molecular features that differentiate PAR from RNA, and offers mechanistic insight into how PAR nucleates phase separation and organizes protein interactions at sites of DNA damage.

## Introduction

Poly(ADP-ribose) (PAR) is a nucleic acid-like homopolymer consisting of repeating adenosine diphosphate ribose (ADP-ribose) units (Fig. 1A), first identified in the seminal work of Chambon and colleagues.^1,2^ PAR is a rapidly synthesized post-translational modification induced in response to DNA damage,^3,4^ where it orchestrates a wide range of cellular processes including DNA repair, chromatin remodeling, RNA biogenesis, protein condensation, intracellular organization, and cell-cycle regulation. ^5–14^ Dysregulation of PAR metabolism is associated with numerous pathological conditions such as neurodegeneration (e.g., Alzheimer’s, Perkinson’s),^15–20^ cancer,^21^ and autoimmune diseases.^22^ Despite PAR’s central importance, the molecular principles underlying both its normal functions and its aberrant behaviors remain unclear. Remarkably, PAR shares striking chemical resemblance to classical nucleic acids, especially polyriboadenylic acid poly-rA (Fig. 1A). This similarity highlights a fundamental biological puzzle: given that RNA is orders of magnitude more abundant in cells (typically ∼ µM) than PAR (typically ∼ nM^23^) and possesses nearly identical chemical building blocks, why does the cell synthesize a distinct polymer, PAR, rather than repurposing RNA to fulfill these same functions?

**Figure 1:**
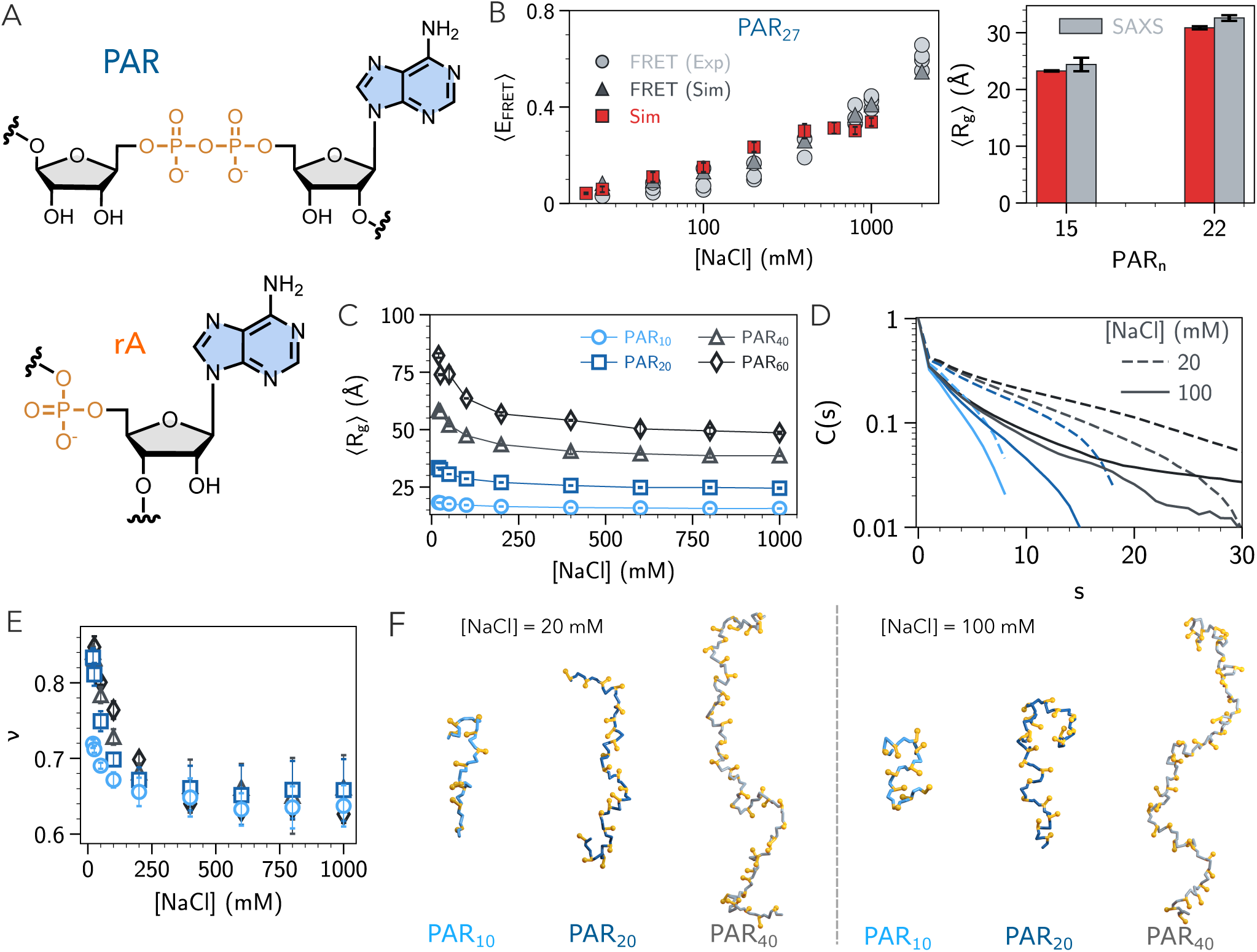
Structural ensemble of PAR: (A) Chemical structure of the monomer unit of PAR (top) and RNA (bottom). (B) (Left) Fluorescence resonance energy transfer (FRET) efficiency for PAR_27_ at 20–1000 mM NaCl from our simulations (red squares) compared with FRET experiments (gray circles) and Monte Carlo simulations^24^ (gray triangles). (Right) Radius of gyration, R*_g_*, for PAR_15_ and PAR_22_ from simulations (red) compared with SAXS.^25^ (C) R*_g_* as a function of monovalent ion concentration and PAR length. (D) Bond-bond autocorrelation C(s) for different PAR length at 20 mM (dashed lines) and 100 mM (solid lines) NaCl. (E) Apparent scaling (Flory) exponent, ν as a function of monovalent ion concentration. (F) Representative snapshots where backbone and bases are highlighted in different colors.

A comprehensive answer to this question is hindered by the limited structural characterization of PAR. Although existing experimental techniques^24–27^ have provided valuable insights into PAR’s structure across different lengths and ionic conditions, they remain challenged by PAR’s intrinsic dynamics and the difficulties associated with its synthesis. Moreover, because PAR contains twice as many phosphate groups as an RNA of equivalent length (Fig. 1A), its conformation is expected to be highly sensitive to ionic conditions. Experimental studies^24,25^ have indeed observed chain-length and ion-dependent structural transitions in PAR, yet these measurements are limited by spatiotemporal resolution. As a result, the microscopic features of PAR structure and the molecular mechanisms by which PAR-ion interactions drive these transitions remain elusive. Computer simulations offer a powerful route to interrogate PAR-ion interactions and capture its structural behavior at atomic detail; however, conventional atomistic simulations face two major limitations: (i) existing force fields are insufficiently accurate for PAR’s unusual chemical topology,^28^ and (ii) the broad, heterogeneous conformational landscape of PAR is difficult to sample within feasible timescales.^25^

To overcome these challenges, we employ a coarse-grained (CG) modeling strategy. We introduce the first CG model that explicitly incorporates ion-PAR interactions using the Reference Interaction Site Model (RISM)^29^ with an implicit solvent representation. This framework reduces the degrees of freedom of both PAR and the surrounding solvent while accurately accounting for ions and their solvation structure, enabling efficient exploration of PAR’s large conformational landscape under diverse ionic environments (Na^+^, Mg^2+^, Ca^2+^). Using this approach, we systematically compare PAR chains of varying lengths, uncovering a striking length-dependent structural transition near physiological ion concentrations: short chains adopt extended conformations, whereas increasing chain length drives a progression from extended to Gaussian coil to compact globular states. These results indicate that PAR’s structural and functional behaviors are governed primarily by its length rather than sequence, distinguishing it from classical nucleic acids. By tuning PAR length, cells may thus generate a continuum of PAR architectures that modulate PAR’s affinity and specificity for distinct DNA-repair and signaling proteins. Moreover, the corresponding changes in ion atmosphere, ion binding, and ion bridging have direct implications^7,13,30^ for PAR’s phase separation propensities, and ultimately for the material properties of the PAR-rich condensates.

Given PAR’s close chemical relationship to nucleic acids, we further compare its structure and ion atmosphere with those of homopolymeric RNAs such as poly-rA and polyribouridylic acid (poly-rU), all sharing the same net charge. Despite carrying the same charge, PAR features a heterogeneous charge distribution (two phosphate groups clustered per monomer) whereas RNA carries a single phosphate per nucleotide. This difference leads to distinct ion sensitivities, ion-phosphate interactions, counterion distributions, and structural transitions, offering insight into why cells preferentially deploy PAR, rather than RNA, for specific biological functions.^9,11^ Our work thus establishes a framework for understanding how PAR serves as a highly charged, multivalent scaffold^30–32^ that can effectively compete with DNA and RNA for binding to nucleic acid-binding proteins central to DNA damage repair and other essential cellular processes.

## Materials and Methods

PAR is a homopolymer resembling RNA, where each unit is composed of two phosphate groups, two riboses, and one adenine base (Fig. 1A). To model PAR, we adapt a similar strategy used for studying RNA folding thermodynamics.^29,33,34^ Each unit is represented by five CG beads: one for each phosphate group, one for each ribose, and one for the adenine base (Fig. S1C). These CG beads are positioned at the center-of-mass (COM) of their respective groups.

The energy function for the FIS model is:

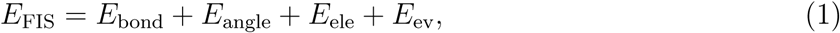

where the first two terms are for bonded and angle interactions, and the last two terms are for electrostatic and excluded volume interactions. We approximate the bonded and angle interactions using harmonic potentials, where the equilibrium values are obtained from the PDB structures of ADP.^35–37^ The spring constants corresponding to the stiffness associated with the bonded and angle constraints are inherited from our RNA model (listed in Table S1). The excluded volume interaction between any two non-bonded beads is computed using the Weeks–Chandler–Anderson (WCA) potential.^38^

The electrostatic interaction between non-bonded phosphate groups, or between any two divalent ions, is computed using the Debye–Hückel screening potential 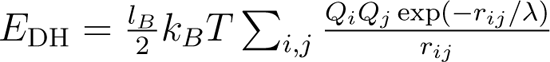 where l*_B_* = e^2^/4πɛ_0_ɛk*_B_*T is the Bjerrum length, and ɛ is the dielectric constant of water. λ = (8πl*_B_*ρ_1_)*^−^*^1*/*2^ is the Debye screening length (ρ_1_ represents the number density of monovalent ions). The effective charge on each phosphate is renormalized to account for counter-ion condensation.^39,40^ In our framework, divalent ions (M^2+^) are explicitly represented in the simulation box, while monovalent ions are treated implicitly. The effective interaction potential between M^2+^ ions and the phosphate group is given by: ^29^

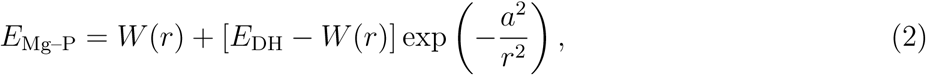

where W (r) denotes the potential of mean force obtained from RISM theory.^41–43^ Following our previous work, ^29^ we set a = 5 Å. The effective potential depends on the temperature as well as the concentrations of monovalent and divalent salts.^44^ The renormalized charge on the phosphate group is determined by solving Eq. S8. A detailed description of the M^2+^–P potential and phosphate charge renormalization can be found in the SI and elsewhere.^29,44–46^

## Results

### Validation of the FIS-PAR model

To validate the model, we perform low-friction Langevin dynamics simulations of PAR_27_ across a wide range of NaCl concentrations (20-1000 mM) at 300 K. We calculate the average Fluorescence Resonance Energy Transfer (FRET) efficiency as 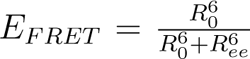, where R*_ee_* is the end-to-end distance of PAR and R_0_ = 59 Å is the Förster radius of the dye used in experiments.^24^ The simulated E*_F RET_* rises as the salt concentration increases, suggesting a structural compaction of PAR at elevated ionic strength. Our results are in quantitative agreement with both experimental measurements and Metropolis Monte-Carlo simulations,^24^ indicating that our model accurately captures the global conformational properties of PAR (Fig. 1B, left). Additionally, the simulated radii of gyration, ⟨R*_g_*⟩, for PAR_15_ and PAR_22_ at 100 mM NaCl are in quantitative agreement with values obtained from small-angle X-ray scattering (SAXS) experiments^25^ (Fig. 1B, right). This further supports the reliability of our model in reproducing the conformational ensemble of PAR across different lengths and ionic conditions.

### PAR undergoes transition from rod to self-avoiding random walk (SAW)-like structure at high monovalent ion concentrations

Given that PAR lengths in cells range from 2 to 200 units of ADP-ribose,^47^ we next investigate how chain length modulates PAR’s structural response to ionic screening. To this end, we perform Langevin dynamics simulations across a broad range of NaCl concentrations (20-1000 mM) for different lengths of PAR (n = 10, 20, 40, 60). For all lengths examined, ⟨R*_g_*⟩ decreases monotonically with increasing salt concentration (Fig. 1C). This compaction reflects electrostatic screening of the negatively charged phosphate groups along the PAR backbone. Notably, the extent of compaction is length-dependent, ranging from 14% for PAR_10_ to 41% for PAR_60_. To further characterize the structural states sampled by PAR, we examine the dimensionless Kratky plot, (qR*_g_*)^2^S(q) vs. qR*_g_*, where S(q) is the backbone structure factor (Eq. S11). The slope at high qR*_g_* (> √3) is used to distinguish between structural states: a positive slope indicates an expanded disordered state, a negative slope suggests a compact globule, and a zero slope corresponds to a Gaussian random coil.^48,49^ In the presence of monovalent ions, PAR remains disordered and expanded (Fig. S3A). To investigate whether salt influences the structural transition in addition to compaction, we analyze the polymeric behavior of PAR using the Flory scaling exponent ν ^50^ (Fig. 1E). For ν values of 1, 0.6, 0.5, and 0.33, the polymer corresponds to a rod, self-avoiding random walk (SAW), Gaussian random coil, and globule structure, respectively.^51^ We extract ν by fitting S(q) versus q in the intermediate q-range using S(q) ∼ q. For all PAR lengths, ν decreases from ≈ 0.7-0.8 to 0.65 as the salt concentration increases, indicating a transition from a rod-like to a SAW-like structure (Fig. 1E). These transitions are consistent with experimental observations.^24,25^

### PAR is devoid of helical structure

The global properties of PAR indicate it is extended and disordered. However, it remains unclear if PAR can adopt a transient helical structure driven by base stacking, as seen in RNA and DNA.^44,52^ To evaluate the possibility of a helical nature, we compute the autocorrelation of bond vectors separated by s bonds, C(s) = ⟨a⃗*_i_* · a⃗*_i_*_+*s*_⟩. Fig. 1D shows that C(s) decays with increasing s and exhibits no periodicity, suggesting that PAR does not adopt any helical conformation^26,27^ (Fig. 1F). This observation is further supported by SAXS data, which show minimal base stacking (1-2 bases) and no helicity in PAR_15_ and PAR_22_ at physiological NaCl concentrations.^25^ The bond correlation decays faster with increasing salt concentration, suggesting of a decrease of the persistent length l*_p_* (relating to C(s) 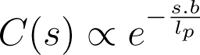, where b is the bond length) which quantifies the chain stiffness. The higher l*_p_* at low salt concentrations is likely due to increased electrostatic repulsion between backbone phosphate charges, which is strongly screened at high salt concentrations. This salt-dependent behavior of l*_p_* in PAR is consistent with FRET experiments.^24^ Interestingly, we find that l*_p_* is not only dependent on salt concentrations but also on PAR length, increasing from approximately 18 Å for PAR_10_ to 55 Å for PAR_60_. Our findings indicate that PAR lacks any transient structure, unlike RNA/DNA, and instead exhibits salt- and length-dependent persistence length.

### PAR undergoes a length-dependent collapse triggered by Mg^2+^

Experiments and simulations have shown that divalent ions, particularly Mg^2+^, induce compaction, folding, and phase separation in PAR^24,25,53–55^ as well as RNA and DNA.^29,56–60^ To assess the effect of Mg^2+^ on PAR, we perform Langevin dynamics simulations for different PAR lengths in a mixture of 20 mM monovalent ion (modeled implicitly) and explicit Mg^2+^ ions ranging from 1 µM to 5 mM. Our simulations show that R*_g_* decreases sharply with increasing Mg^2+^ concentration in a sigmoidal manner for all PAR lengths, consistent with a cooperative response (Fig. 2A). Interestingly, the transition is notably steeper for longer PARs, suggesting that cooperativity increases with PAR length. Importantly, all PAR chains undergo compaction around the same narrow range of Mg^2+^concentration (∼0.1 mM) (Fig. 2A) despite their varying net charges.

**Figure 2:**
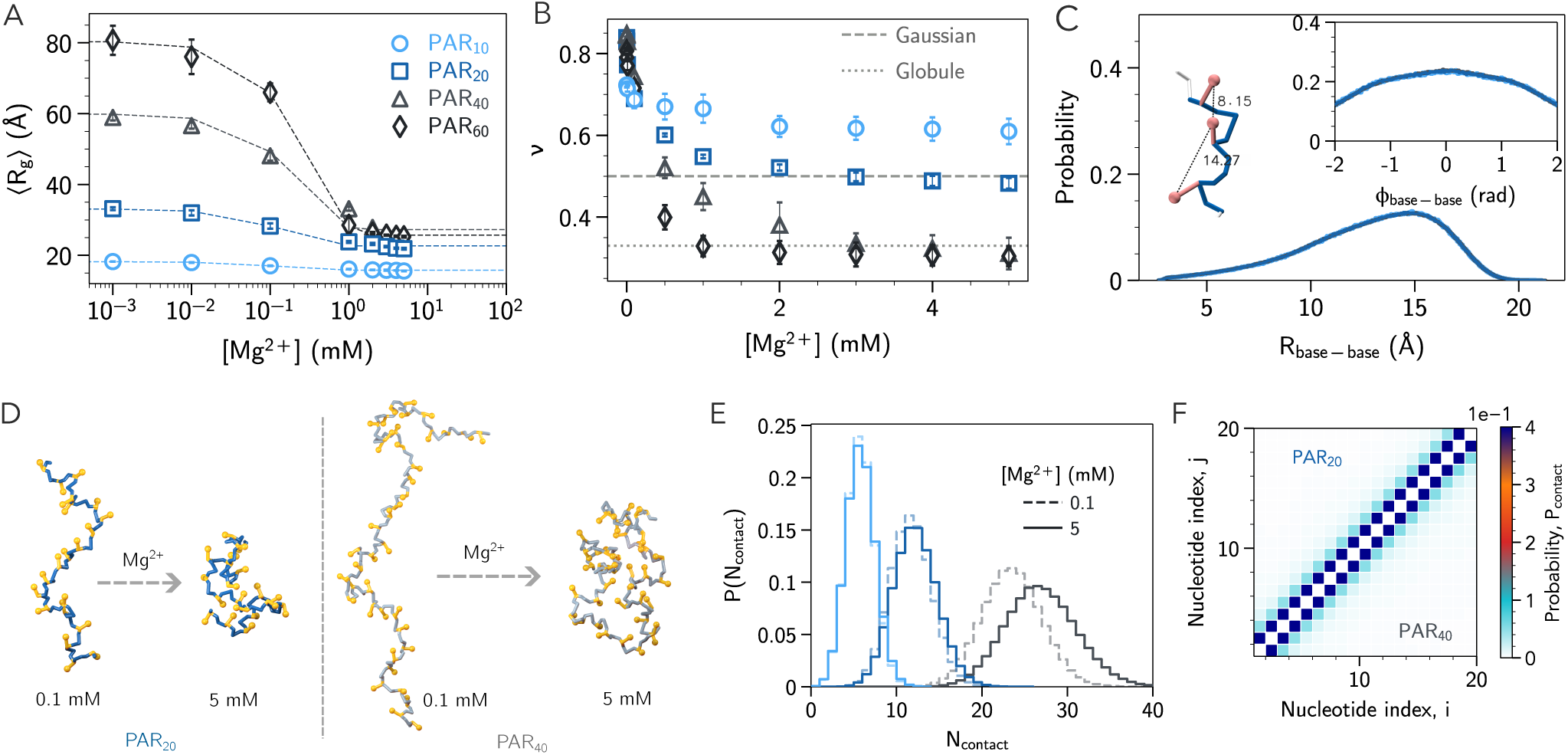
Length-dependent collapse of PAR in the presence of Mg^2+^: (A) Radius of gyration at various Mg^2+^ concentrations for different lengths of PAR (with 20 mM NaCl in the background). (B) Apparent scaling exponent ν at different Mg^2+^ concentrations. The two horizontal lines correspond to the Gaussian random coil (dashed) and compact globule (dotted). (C) Probability distribution of consecutive base-base distance and dihedral angle (inset) of PAR. Also shown is a segment of PAR_20_ highlighting unstacked bases. (D) Representative snapshots show compaction of PAR at elevated Mg^2+^ concentrations. (E) Number of intramolecular contacts at 0.1 mM (dashed lines) and 5 mM (solid lines) Mg^2+^ concentrations for three different PAR lengths. (F) Contact probability between two units in the same chain in 1 mM Mg^2+^ for PAR_20_ (upper diagonal) and PAR_40_ (lower diagonal, only shown 1–20). For PAR_40_, the remaining units (21–40) follow the same pattern.

The Flory exponent ν further validates the length-dependent structural transition of PAR in Mg^2+^ (Fig. 2B). At low Mg^2+^, all chain lengths remain in extended states (ν > 0.7). However, at high-ion concentrations, PAR structure becomes length dependent. For PAR_10_, ν decreases to 0.6, indicating the transition from a rod-like to SAW-like structure. In contrast, medium sized chains such as PAR_20_ behave as Gaussian random coils (ν ≈ 0.5), while longer PAR_40_ and PAR_60_ even fold up to globular structures at elevated Mg^2+^ conditions (Fig. 2B,D). The Kratky plots show the same trend (Fig. S3B). As a result, the degree of collapse becomes highly dependent on PAR length. The greater compaction in longer chains observed here is in line with experiments^25^ and likely arises from intrinsic chain stiffness.^61^ In summary, our result suggests that PAR compaction is strongly influenced by ion concentration, valency, and chain length.

Collapse of nucleic acids is typically driven by base stacking interactions. ^44,52,62^ However, due to its lengthening backbone compared to RNA, the bases in PAR are relatively far away from each other, which weakens stacking interactions. To explore the origin of PAR collapse, we first check the potential for base stacking by calculating the relative distance and orientation between consecutive bases in PAR. Fig. 2C shows the consecutive bases indeed remain far away from each other (R_base-base_ is primarily between 10-20 Å), and are randomly oriented (almost uniform angle distribution). This indicates that PAR forms minimal base stacking, and thus their contributions to PAR collapse at high ion concentrations are minor. In addition, the contact map (Fig. 2F) shows that most contacts are along the diagonal, local and uniform, suggesting these contacts are non-specific and Mg^2+^-dependent (Fig. 2E).

### Comprehensive mapping of PAR ion atmosphere

Since the collapse of PAR is not driven by base stacking and highly dependent on Mg^2+^ concentration, we posit that ion-mediated interactions likely promote these contact formation, ultimately leading to PAR compaction. To quantify ion-mediated interactions in PAR, we calculate the radial distribution function, g*_Mg−P_*, between Mg^2+^ and phosphate beads. g*_Mg−P_* exhibits two peaks at r ≈ 2.5 and 5.0 Å (Fig. 3A), indicating two distinct binding modes: direct ion-phosphate interactions (inner-sphere, IS) and water-mediated interactions (outer-sphere, OS)^29,45,63^ (illustrated in Fig. 3B). Notably, the populations of both IS and OS bindings increase with PAR length, indicating a stronger ion binding affinity for longer chains (Fig. 3A). As a result, the number of excess ions Γ*_Mg_*2+, which quantifies how many ions are attracted towards the chain, ^64^ is consistently higher for longer chains (Fig. S4). Additionally, Γ*_Mg_*2+ increases at high Mg^2+^ concentrations for all PAR lengths. Thus, both g*_Mg−P_*and Γ*_Mg_*2+ confirm that longer PAR chains attract more ions, likely contributing to their higher degree of collapse. Strikingly, the ion concentration converges to bulk values (g*_Mg−P_* (r) → 1) at larger distances for longer PARs (Fig. 3A, inset), suggesting that the ion atmosphere of longer chain is more extended.

**Figure 3:**
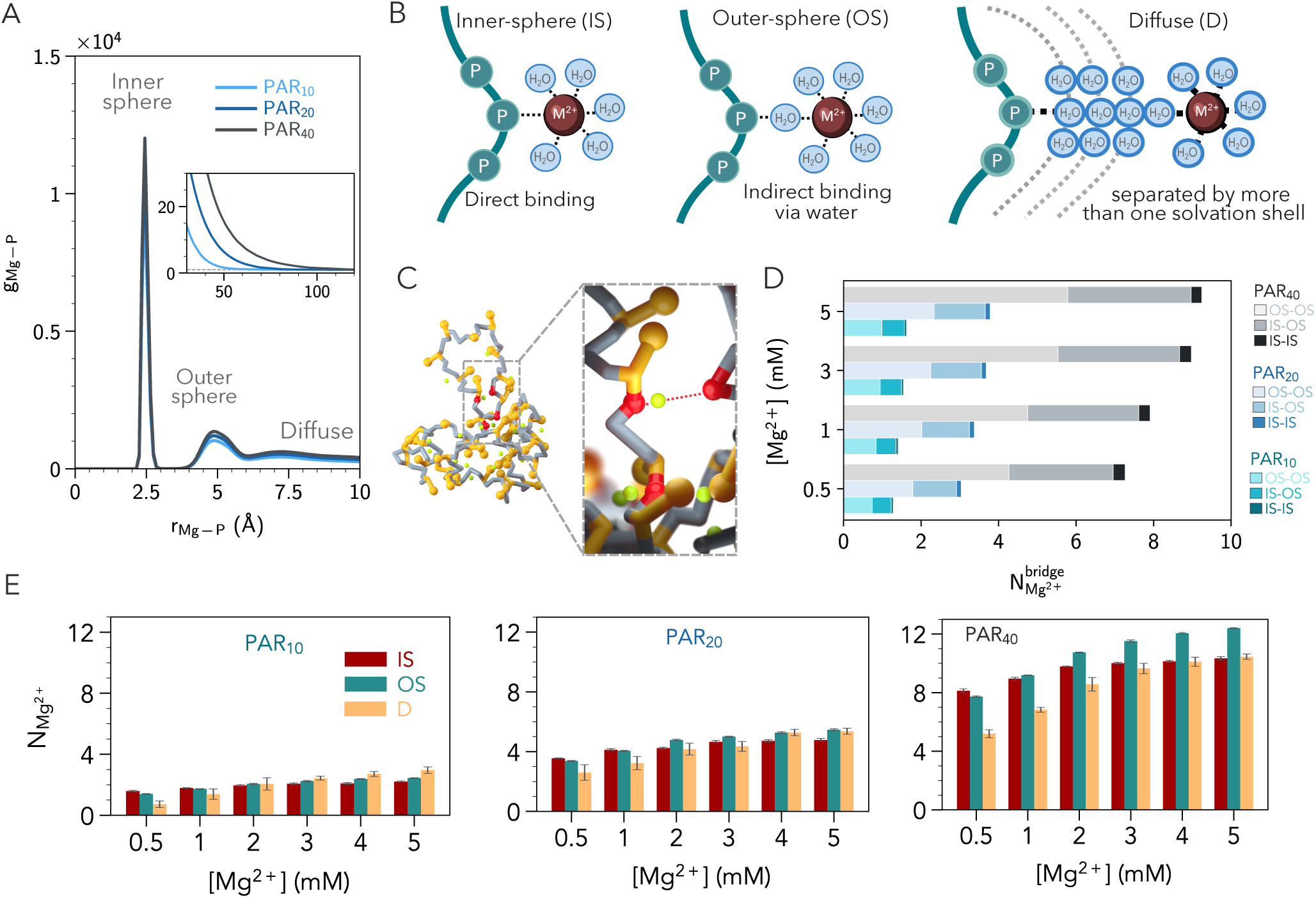
Ion atmosphere around PAR: (A) Radial distribution function, g*_Mg−P_*, between Mg^2+^ ions and phosphate for various PARs with different lengths. The inset shows the decay of g*_Mg−P_* at large separation. (B) Schematics of ion binding modes: inner-shell (IS), outer-shell (OS), and diffuse (D). Phosphate backbone, divalent ion (M^2+^) and water are shown in teal, brown and cyan, respectively. Interactions between ions and phosphate are highlighted by dotted lines. (C) A snapshot highlights bridging ions (lime) and phosphate beads (red). (D) Number of ions for different modes of bridging. (E) Partition of ion atmosphere into each binding mode for various PARs at different Mg^2+^ concentrations.

In addition to these IS/OS ions shown in Fig. 3A, a significant population of diffusely interacting ions exists, which contributes to the long range decay of g(r) and is often referred to as diffuse ions (Fig. 3B, right). As in previous work, ^45^ we partition the PAR ion atmosphere into IS (0 < r*_Mg−P_* < 3.2 Å), OS (3.2 ≤ r*_Mg−P_* < 6.1 Å) and diffuse ions (Fig. 3E). The number of diffuse ions are then calculated as N*_Mg_*_2+_ = Γ*_Mg_*2+ − N*_Mg_*_2+_ − N*_Mg_*_2+_. At low Mg^2+^ concentrations (0.5–1 mM), ions mostly bind in the IS and OS regions across all PAR lengths (Fig. 3E), suggesting a compact ion atmosphere. At higher ion concentrations (2–5 mM), the ion binding pattern becomes length-dependent. For short PAR_10_, diffuse ions now contribute the most (D > OS > IS), supporting the idea that the ion atmosphere expands at higher-ion concentrations. Remarkably, longer chains such as PAR_40_ still show a slight preference for OS ions, indicating a smaller shift from IS to OS (instead of diffuse) as PAR length increases, reflecting a stronger affinity for bound ions in longer chains.

Due to a large number of bound ions in IS and OS, some of them could bind simultaneously to two (or more) phosphate groups, acting as bridges and thus facilitating secondary and tertiary interactions, leading to the collapse (Fig. 3C-D). Such bridging ions have been found to play important roles in folding and function of many nucleic acids such as ribozymes.^29,34,45,56,60,65^ Indeed, we find that the number of bridging ions increases with chain length (Fig. 3D), with PAR_20_ having roughly twice the bridging ions of PAR_10_ while PAR_40_ having 7–8 times more. This trend is observed across all bridging modes (IS-IS, IS-OS, OS-OS). In PAR_10_, where the number of bridging ions is low, the chain remains extended (Fig. 2B). By contrast, higher numbers of bridging ions in PAR_20_ and PAR_40_ drive structural transitions to Gaussian and globular states (Fig. 3D), reinforcing that ion bridging drives structural collapse of PAR.

### PAR collapse is distinct from RNA

Although PAR shares the same chemical building blocks as poly-rA (Fig. 1A), and poly-rA is abundant in cells,^66^ nature selectively uses PAR for a wide range of specialized functions.^13,20,32,67^ To understand why, we compare the structural properties of PAR with poly-rA and poly-rU. Given that PAR contains twice the number of phosphate groups (Fig. 1A), we select rU_40_ and rA_40_ to match the phosphate count in PAR_20_.

Similar to PAR, both rU_40_ and rA_40_ collapse at high Mg^2+^ concentrations (Fig. 4A).^44,52^ However, the extent of compaction is much greater for PAR, as the change in R*_g_* drops from 34% for PAR to 22% for the two RNAs. Moreover, the transition is steeper for PAR, indicating a higher degree of cooperativity relative to RNA. Importantly, PAR collapses at a much lower critical Mg^2+^ concentration than RNA (0.1 mM vs. 1 mM), highlighting PAR’s increased sensitivity to divalent ions (Fig. 4A). The Kratky plots further demonstrate the sharp structural difference between PAR and RNA at high Mg^2+^ concentrations (Fig. S5). Even at 5 mM Mg^2+^, both RNA sequences remain in an unfolded/extended state. In contrast, PAR adopts a Gaussian/globule-like structure at the same conditions. Similarly, the Flory exponent reveals that PAR collapses to a Gaussian random coil (ν ≈ 0.5),^26,28^ while the two RNAs can only reach a SAW-like state (ν ≈ 0.6) at high Mg^2+^ concentrations (Fig. 4B-C). This suggests a more pronounced structural transition in PAR compared to RNA. Contrary to PAR, which shows minimal base-stacking, both RNAs populate stacked conformations where two consecutive bases are close and aligned to each other (Fig. S6) and thus display transient helical conformations, especially for rA_40_ (Fig. S7).

**Figure 4:**
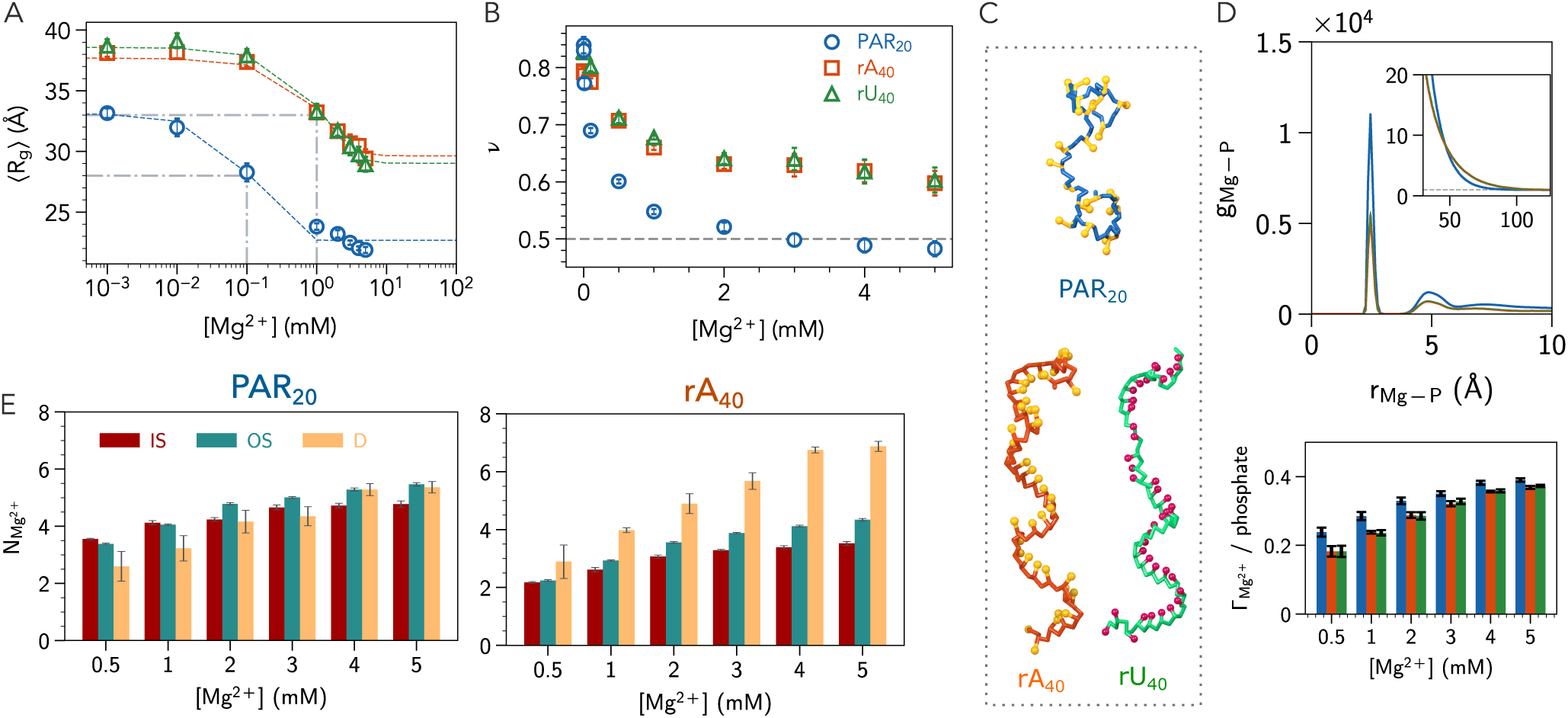
Distinct Structural Transitions and Ion Atmospheres in PAR versus RNA: (A) Radius of gyration of PAR_20_ (blue), rA_40_ (orange) and rU_40_ (green) at various Mg^2+^ concentrations (in the background of 20 mM NaCl). The dashed lines represent the sigmoidal fit. The gray lines correspond to the critical Mg^2+^ concentrations for the transition. (B) Apparent scaling exponent for the three molecules. (C) Representative snapshots from simulations where backbone and bases are in different colors. (D) (top) Radial distribution function, g*_Mg−P_*, between Mg^2+^ ions and phosphate beads for the three molecules. The inset shows the decay of g*_Mg−P_* at large separation. (bottom) Number of excess Mg^2+^, Γ*_Mg_*2+, per phosphate at different Mg conditions. (E) Number of inner-shell (IS), outer-shell (OS), and diffuse (D) ions at different Mg^2+^ concentrations for (left) PAR_20_ and (right) rA_40_.

Remarkably, the structural differences between PAR and RNA are highly dependent on ion valency (Na^+^ vs. Mg^2+^). While PAR remains more extended than RNA in monovalent ions (Fig. S8), it collapses to a much more compact state under the presence of Mg^2+^ (Fig. 4A). This contrasting behavior between PAR and RNA with respect to ion valency aligns with experimental findings showing that PAR requires lower Mg^2+^ concentration but higher monovalent ion concentration than RNA to achieve the same degree of collapse.^24^

### PAR attracts more ions than RNAs with distinct ion atmosphere

Due to its much smaller size compared to RNA, the ion atmosphere of PAR is also highly compact. This can be seen by significantly higher peaks for both IS and OS regions compared to RNA (Fig. 4D, top), suggesting that PAR attracts more ions. As a result, Γ*_Mg_*2+ per phosphate for PAR is consistently higher than RNAs (Fig. 4D, bottom). This behavior can be attributed to the two phosphate groups next to each other, boosting its charge density compared to RNA.

Strikingly, despite attracting more ions, the overall ion atmosphere of PAR is more compact than RNA, with a crossover between them occurring at r ≈ 45 Å (Fig. 4D, inset). The local Mg^2+^ concentration around PAR decays to bulk values (g*_Mg−P_* (r) → 1) more rapidly than around RNA, indicating that ions are confined closer to the PAR backbone. PAR achieves this primarily through enhanced IS and OS binding, with diffuse ions playing a minor role except at high concentrations (Fig. 4E, left). In contrast, for RNA sequences, Mg^2+^ predominantly interacts via diffuse ions, consistent with previous studies^29,45,46^ (Fig. 4E, right and Fig. S10C, right). Moreover, the number of IS and OS ions is significantly higher for PAR compared to RNA, emphasizing the role of sequence in shaping the ion atmosphere. Our previous simulations ^46^ have shown that the extent of the RNA ion atmosphere depends strongly on its structural complexity, with disordered RNAs exhibiting a more expanded ion atmosphere than folded RNAs. In this regard, PAR’s ion atmosphere resembles that of folded RNAs, even though PAR itself is structurally disordered. We attribute this behavior to the different charge density of PAR, which enhances electrostatic attraction and draws counterions closer to the chain compared to disordered RNAs.

### Ion identity affects PAR and RNA differently

We next examine how monovalent ion concentration (from 20–100 mM) and divalent ion identity (Mg^2+^ vs. Ca^2+^) modulate the ion atmosphere of PAR and RNA. As monovalent ion concentration increases, Γ*_Mg_*2+ decreases for both molecules due to enhanced screening of phosphate charges (Fig. S9B, left). In RNA, this leads to reduced IS binding (Fig. S9A, left). In contrast, PAR shows the opposite behavior: despite a lower Γ*_Mg_*2+, IS binding increases at higher monovalent ion concentrations. This 1–2 fold increase is coupled with a twofold reduction in diffuse ions (Fig. S10B, left), reflecting a redistribution from diffuse to IS binding. This shift arises from a modest collapse induced by monovalent ions (≈15%; Figs. S11, S12A), which enhances nucleotide accessibility for ion bridging and increases the fraction of bound ions. By comparison, RNA shows a uniform decrease in all ion populations (IS, OS, D) at elevated Na^+^ concentrations (Fig. S10B, middle-right). Thus, while IS binding increases in PAR but decreases slightly in RNA, these opposite responses reflect intrinsic differences in size and charge density: PAR, being smaller and more highly charged, requires a larger number of bound ions for charge neutralization (Fig. S12A), whereas the larger and less charge-dense RNA requires fewer.

Switching the divalent ion from Mg^2+^ to Ca^2+^ leaves the overall ion binding, Γ, unchanged (Fig. S9B, right). However, Ca^2+^ displays a stronger preference for IS over OS binding in both PAR and RNA (Fig. S10D). Unlike monovalent ion, this enhancement in IS binding occurs without structural collapse (Fig. S12B). Instead, it stems from the lower free energy barrier^29,45,46,56^ for the OS-to-IS transition in Ca^2+^ relative to Mg^2+^. Ca^2+^’s larger size and lower charge density facilitate water release from its first solvation shell, making direct phosphate coordination more favorable.

### Ion-mediated long-ranged bridging underlies PAR collapse

We suspected that the preference for IS/OS binding in PAR, as opposed to diffuse binding in RNA, is due to the ion bridging. We indeed observe that PAR has 4–6 times more bridging ions than RNA (Fig. 5B, left), indicating that despite carrying the same net charge, PAR’s blocky charge pattern promotes bridging more effectively than RNA. Bridging ions are categorized as IS–IS, IS–OS, or OS–OS based on ion-phosphate distances (Fig. 5A). Across all Mg^2+^ concentrations and both sequences, OS–OS bridging is most abundant, followed by IS–OS and IS–IS (Fig. 5B, left). Because IS–IS bridging requires a single ion discarding two hydration water molecules, it is enthalpically unfavorable. In addition, bringing two phosphate groups into close proximity to the ion (≲ 3.2 Å) incurs an entropic penalty, making IS–IS bridging less favorable than IS–OS and OS–OS modes. We also observe a small fraction of ions that bridge more than two phosphate groups in PAR, whereas such higher-order bridging is negligible in RNA (Figs. S13, S14).

**Figure 5:**
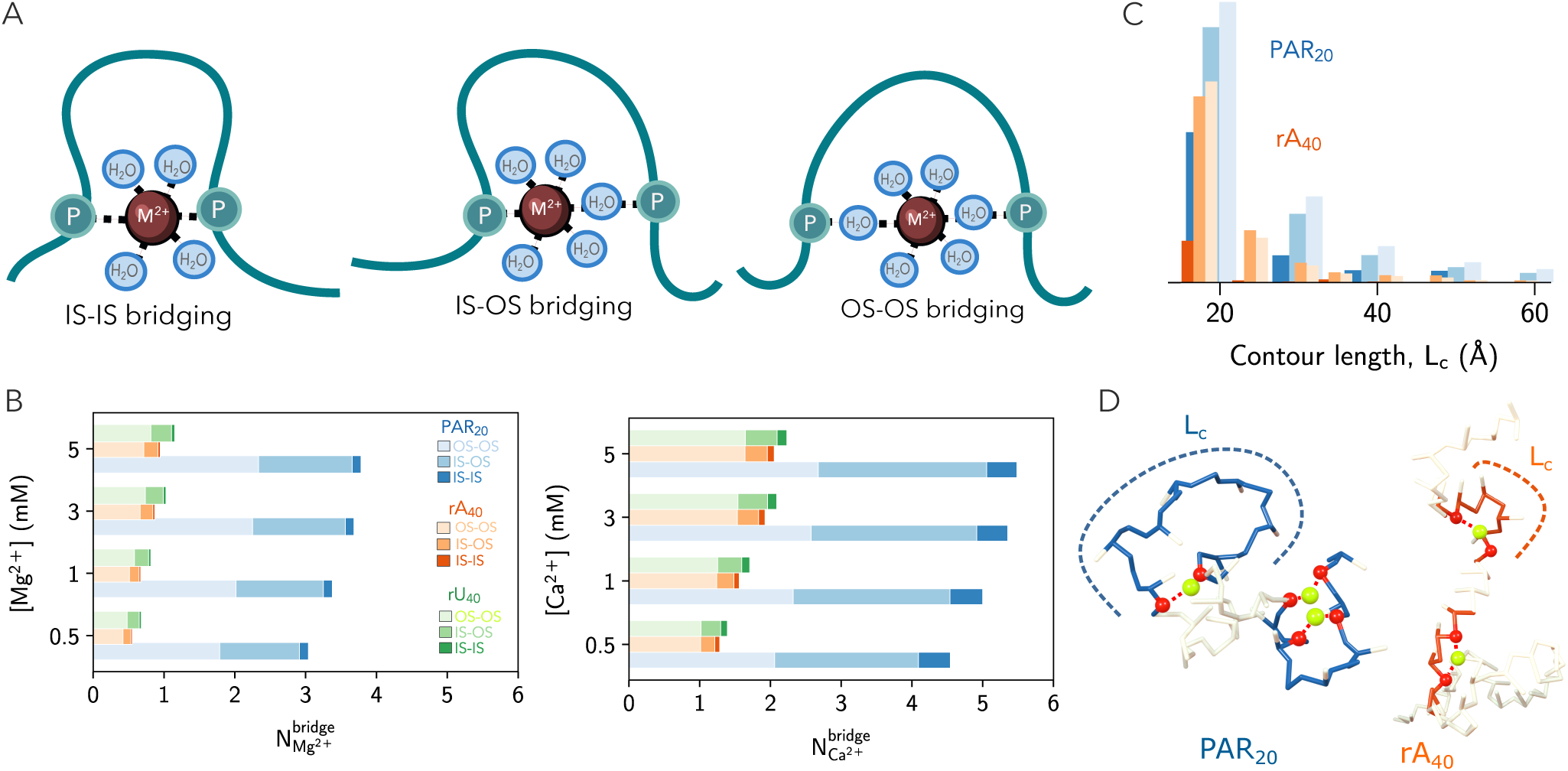
Ion-mediated bridging induces PAR collapse: (A) Schematics of ion bridging: IS-IS, IS-OS and OS-OS. Phosphate backbone, divalent ion (M^2+^) and water molecules are shown in teal, brown and cyan color. (B) Average number of bridging ions in a mixture of 20 mM NaCl + (left) Mg^2+^ or (right) Ca^2+^ for PAR_20_ (blue), rA_40_ (orange), and rU_40_ (green). (C) Probability of long-range bridging for a contour length separation, L*_c_*, for various modes. (D) Representative snapshots illustrating contour distances L*_c_*, highlighted in blue for PAR_20_ and orange for rA_40_. Phosphate beads, Mg^2+^ ions, and their coordination are represented by red spheres, lime spheres, and red dashed lines, respectively.

Not only does PAR have a larger number of bridging ions, but these ions also bridge phosphate groups over greater contour distances compared to RNA. To quantify this, we calculate the contour separation (L*_c_*) between bridged phosphate pairs (Fig. 5C). In PAR, Mg^2+^ can bridge phosphate groups separated by more than 60 Å, whereas bridging in RNA is limited to ≈ 40 Å (Figs. 5C, D).

For PAR, the populations of long-range bridging are ≈15%, 27%, and 32% for IS-IS, IS-OS, and OS-OS modes, respectively, while the corresponding RNA values are substantially lower (≈5%, 23%, 22%). These results demonstrate that Mg^2+^-mediated long-range bridging is significantly more favorable in PAR, owing to its blocky charge distribution, and that these interactions strongly facilitate chain collapse. Increasing number of bridging ions progressively shift the R*_g_* distribution toward smaller values, confirming their central role in driving PAR collapse (Fig. S15). RNA, on the other hand, accommodates far fewer bridging ions overall(Fig. S16), resulting in a substantially weaker global collapse (Fig. 4A).

Replacing Mg^2+^ with Ca^2+^ increases the total number of bridging ions for both PAR and RNAs (Fig. 5B). For PAR, this increase arises mainly from enhanced IS–IS and IS–OS bridging, while OS–OS bridging remains largely unchanged. Strikingly, the opposite trend is observed for two RNAs, where the increase in bridging occurs predominantly through OS–OS bridging. Despite these differences in the magnitude and mode of bridging, the overall bridging pattern remains nonspecific and uniformly distributed along the backbone across all examined conditions (Fig. S17-S19).

## Discussion and Conclusions

PAR, often attached to proteins as a post-translational modification, is widely available in the cell,^68^ including the nucleus, cytoplasm, and even extracellular matrix. It acts as a “friend in need,” assisting cellular recovery from stress through the phase separation of DNA/RNA-binding proteins. In this study, we introduce the first accurate CG model to investigate the structure and ion atmosphere of PAR under diverse ionic conditions. Our simulations provide molecular insights into PAR’s structural transitions and enable direct comparison with RNA, shedding light on two key questions: (1) How can PAR, present only at nM concentrations, effectively compete with µM RNA to drive protein phase separation, and (2) Why does the cell synthesize PAR specifically for certain functions rather than repurposing abundant RNA?

We find that PAR undergoes a sharp structural collapse at ion concentrations nearly ten-fold lower than RNA, demonstrating its greater sensitivity to multivalent cations. Functionally, PAR facilitates protein condensation by acting as a multivalent, highly negatively charged scaffold that strongly engages intrinsically disordered proteins and regions (IDPs/IDRs) through electrostatic interactions.^31^ This increased sensitivity allows PAR to trigger phase separation at far lower concentrations than RNA. Consistent with this, cellular PAR level (∼nM)^23^ is substantially lower than RNA (∼ µM), yet PAR is sufficient to induce phase separation: nM PAR drives FUS droplet formation, whereas µM RNA is required under comparable conditions.^30^ Moreover, PAR can compete with DNA/RNA for binding to nucleic acid-binding proteins. Due to its strong response to multivalent ions, PAR can displace DNA/RNA and recruit proteins that are normally bound to these nucleic acids, thereby facilitating its functions. For example, PAR interacts with DNA-bound histones, disrupting histone-DNA contacts and rendering damaged DNA accessible for repair. ^32^

Our work reveals that RNA’s ion atmosphere is far more extended, facilitating long-range interactions that may recruit distant or non-essential proteins. In contrast, PAR’s compact ion atmosphere restricts interactions to nearby, functionally relevant targets, providing a mechanistic explanation for its selective deployment despite its much lower abundance. Interestingly, we also demonstrate that Ca^2+^ exhibits a stronger preference for binding to PAR via the IS coordination. Because IS-bound ions directly contact with phosphate groups, these interactions are substantially stronger than those of OS or diffusive ions. Consequently, we surmise that PAR will display a higher affinity for Ca^2+^ in a mixed environment. This prediction aligns with in vivo and in vitro observations showing that PAR preferentially recruits Ca^2+^ to promote biomineralization in the bone extracellular matrix, especially around collagen fibrils where PAR facilitates the formation of spheroid mineral deposits.^69,70^ Such mineralization does not occur with canonical nucleic acids like DNA or RNA, consistent with our finding that RNA exhibits much weaker IS binding. Beyond mineralization, the stronger IS preference of Ca^2+^ also enhances PAR’s phase separation propensity: in vitro, PAR forms larger liquid-like droplets at much lower Ca^2+^ concentrations than Mg^2+^,^54,55^ similar to trends observed for RNA.^71^

Our analyses across PAR chain lengths reveal that Mg^2+^ induces different degrees of collapse depending on polymer length. At the same Mg^2+^ concentration, PAR_10_ remains extended, PAR_20_ behaves as a random coil, whereas longer PAR_40_/PAR_60_ adopt globular structures. This chain length-dependence is not observed in RNA or DNA, whose structural responses are governed primarily by sequence.^44^ Interestingly, no cellular analogs of PAR with alternative nucleotide compositions are known (such as poly(GDP), poly(CDP), or poly(UDP)). Based on our findings, we propose that PAR tunes its function primarily through variations in chain length. This hypothesis is supported by experimental evidence showing that PAR’s binding specificity depends strongly on its length.^14,47,72–77^ Recent work^78^ shows that short PAR_5_ exhibits strong binding preferences for specific DNA- and RNA-binding proteins, while longer PAR_20_-PAR_50_ prefers distinct classes. We also demonstrate that bridging ions in PAR increases with chain length. Because ion-mediated bridging is a major driver of nucleic acid phase separation,^58^ longer PAR chains should exhibit a higher propensity for phase separation. Experiments support this view:^30^ monomeric and dimeric PAR are unable to induce condensation. PAR_5_ is the minimum length required, and condensate formation becomes increasingly pronounced with longer chains. This length dependence also influences condensate dynamics and material properties: shorter chains form dynamic, liquid-like droplets, whereas longer chains form more solid-like aggregates.^30^

Beyond mechanistic insight, our model provides a tractable framework to explore PAR-protein interactions and the competition between PAR and RNA in future. Collectively, these results provide a blueprint for designing nucleic acid sequences with tailored charge blockiness for novel biomaterials, including ion sensors and artificial ion-responsive systems.

## Acknowledgment

We gratefully acknowledge financial support from the University at Buffalo to H.T.N.. Computational resources were provided by the Center for Computational Research at the University at Buffalo.

## TOC Graphic

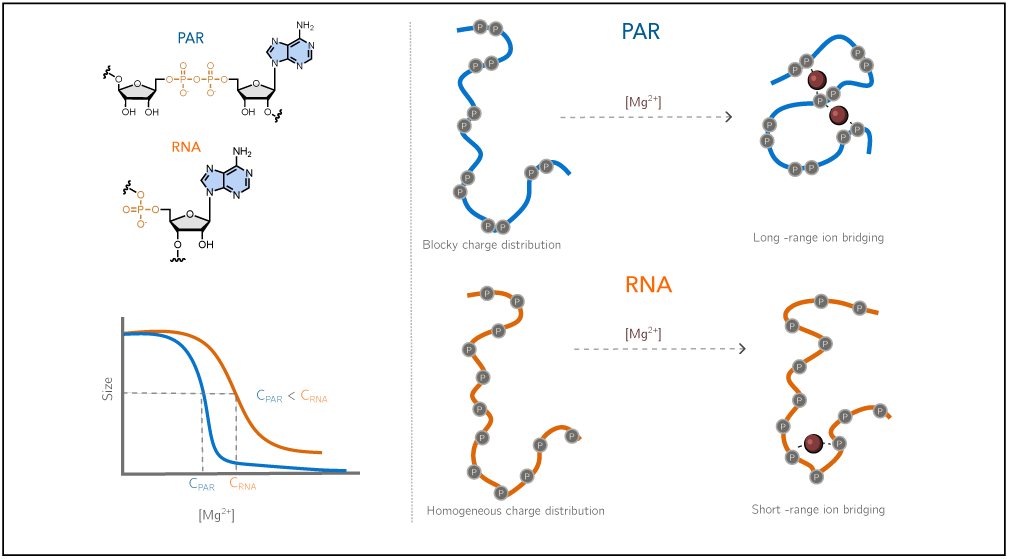

