## Supplementary File for "Why PAR is not RNA: ion atmospheres, bridging interactions, and ion-induced structural collapse"

### Five-Interaction-Site (FIS) Model for Poly(ADP-Ribose):

Poly(ADP-Ribose) (PAR) is a homopolymer resembling RNA, where each unit is composed of two phosphate groups, two riboses, and one adenine base (Fig. S1). To model PAR, we adapt a similar coarse-grained (CG) strategy used for studying RNA folding thermodynamics.[1–3] Each unit is represented by five CG beads: one for each phosphate group, one for each ribose, and one for the adenine base (Fig. S1). These CG beads are positioned at the center-of-mass (COM) of their respective groups.

The energy function for the FIS model is:

$$E_{FIS} = E_{bond} + E_{angle} + E_{ele} + E_{ev}, \tag{S1}$$

where $E_{bond}$, $E_{angle}$, $E_{ele}$ and $E_{ev}$ are the energy terms for bonded, angle, electrostatic and excluded volume interactions, respectively. The bonded and angle interactions are approximated using harmonic potentials:

$$E_{\text{bond}} = k_r (r - r_0)^2, \tag{S2}$$

$$E_{\text{angle}} = k_\theta (\theta - \theta_0)^2, \tag{S3}$$

where the equilibrium values, $r_0$ and $\theta_0$, are obtained from the PDB structures of ADP.[4–6] The spring constants, $k_r$ and $k_\theta$, correspond to the stiffness associated with the bonded and angle constraints, inherited from our RNA model. The specific values of these parameters are listed in Table S1.

The excluded volume interaction ($E_{\text{ev}}$) between any two non-bonded beads is computed using the Weeks–Chandler–Anderson (WCA) potential:[7]

$$E_{\text{ev}} = \begin{cases} \epsilon \left[ \left(\frac{\sigma_{ij}}{r_{ij}}\right)^{12} - 2\left(\frac{\sigma_{ij}}{r_{ij}}\right)^{6} + 1 \right] & \text{for } r \leq \sigma_{ij} \\ 0 & \text{for } r > \sigma_{ij}, \end{cases} \tag{S4}$$

where $\epsilon = 1$ kcal/mol is the strength of the interaction, and $\sigma_{ij} = R_i + R_j$, where $R_i$ and $R_j$ are the radii of the CG beads (listed in Table S1).

The electrostatic interaction is partitioned into two separate contributions. First, the electrostatic interactions between nonbonded phosphate beads, or between divalent ions, is computed using the Debye–Hückel screening potential:

$$E_{ele1} = \frac{l_B}{2} k_B T \sum_{i,j} \frac{Q_i Q_j \exp(-\frac{r_{ij}}{\lambda})}{r_{ij}}, \tag{S5}$$

where $l_B = \frac{e^2}{4\pi\epsilon_0\epsilon_r k_B T}$ is the Bjerrum length, and $\epsilon$ is the dielectric constant of water. $\lambda = (8\pi l_B \rho_1)^{-1/2}$ is the Debye screening length ($\rho_1$ represents the number density of monovalent ions). The charge of divalent ions is $Q_M = 2$, while the effective charge of phosphate $Q_P$ is renormalized to account for counter-ion condensation. In the absence of divalent ions, $Q_P = \frac{b}{l_B}$, where $b = 4.4$ Å is the average spacing between two phosphate beads. The effect of $b$ on $Q_P$ is shown in Fig. S2. In the presence of divalent ions, we solve the following equations for $Q_P$:

$$\theta_1 + 2\theta_2 = \theta = 1 - \frac{b}{l_B}, \tag{S6}$$

$$\frac{e\theta_2}{C_2 V_2} = 2 \ln \left( \frac{e\theta_1}{C_1 V_1} \right) \tag{S7}$$

$$V_i = 4\pi e b^3 (1 + Z_i) \left( \xi - \frac{1}{Z_i} \right) \tag{S8}$$

where $\theta_1$ and $\theta_2$ represent the numbers (per phosphate) of condensed monovalent and divalent ions, respectively. $C_1$, $C_2$, $V_1$, and $V_2$ denote the bulk concentration and effective condensation volume of monovalent and divalent ions, respectively. Since monovalent ions are treated implicitly, the effective charge on the phosphate group, considering only monovalent ion condensation, is given by $Q_P = 1 - \theta_1$.

The second part of $E_{ele}$ is the effective potential between $M^{2+}$ and phosphate, calculated using:[2]

$$E_{\text{Mg–P}} = W(r) + [E_{\text{DH}} - W(r)] \exp\left(-\frac{a^2}{r^2}\right), \tag{S9}$$

where $W(r)$ represents the potential of mean force computed using RISM theory, and $E_{\text{DH}}$ is the Debye–Hückel potential between $M^{2+}$ and P screened by monovalent salt ions (Eq. S5). We set $a = 5$ Å as in previous work.[2] The effective potential is thus dependent on temperature, as well as the concentrations of both monovalent and divalent salts. The total electrostatic is thus $E_{ele} = E_{ele1} + E_{\text{Mg-P}}$.

### Simulation details

In our simulations, divalent ions ($M^{2+}$) are explicitly represented in a periodic cubic box, while monovalent ions are treated implicitly. The initial PAR structure is generated as a linear chain oriented along the Z-axis, and divalent ions are randomly distributed within the box. The number of divalent ions is fixed at $N_{M^{2+}} = 400$, while the box length is adjusted to achieve the desired ion concentration. For RNA simulations, we use an updated version of the TIS model.[8]

We perform low-friction Langevin dynamics simulations at temperature of $T = 300\,\text{K}$ using the LangevinIntegrator module in OpenMM.[9] The equation of motion for Langevin dynamics is given by:

$$m_i \frac{\mathrm{d}\mathbf{v_i}}{\mathrm{d}t} = \mathbf{f_i} - \gamma m_i \mathbf{v_i} + \mathbf{R_i}, \tag{S10}$$

where $m_i$ and $\mathbf{v_i}$ are the mass and velocity of particle $i$, respectively, $\mathbf{f_i}$ is the deterministic force exerted by other particles on particle $i$, and $\mathbf{R_i}$ is an uncorrelated random force on particle $i$, with components sampled from a normal distribution with mean zero and unit variance. The friction coefficient $\gamma$ is set to $1\,\text{ps}^{-1}$. We integrate the Langevin equation using a time step of $\delta t = 2\,\text{fs}$.

### Data Analysis

**Structure factor:** The structure factor is computed as:

$$S(q) = \frac{1}{N_s^2} \sum_{i=1}^{N_s} \sum_{j=1}^{N_s} \frac{\sin(q r_{ij})}{q r_{ij}}, \tag{S11}$$

where $q$ is the wave vector and $r_{ij}$ is the distance between the sugar beads $i$ and $j$. $N_s$ represents the number of sugar beads attached to the base. To ensure consistency between PAR and RNA,

$S(q)$ is computed using only the sugar beads that are attached to the base.

**Local ion concentration:** The local divalent ion concentration, $C_{M^{2+}}$ (M = Mg, Ca) is computed for each nucleotide considering phosphate beads of PAR/RNA using the equation,

$$C_{M^{2+}} = \frac{1}{V_c}\int_0^{r_c} 4\pi r^2 \rho(r) dr, \tag{S12}$$

where $V_c$ is the spherical volume of cutoff radius $r_c$. $\rho(r)$ is the density of divalent ions at a distance $r$ from the phosphate. The cutoff distance for IS $Mg^{2+}$ and $Ca^{2+}$ ions are $0 < r_c < 3.2$ and $0 < r_c < 3.7$ Å, respectively. The same for OS $Mg^{2+}$ and $Ca^{2+}$ ions are $3.2 \leq r_c < 6.1$ and $3.7 \leq r_c < 6.6$ Å.

**Table S1: Parameters for the FIS-PAR model.** P, S, and B represent the phosphate, sugar, and base, respectively. $m$ and $R$ are the bead mass and radius. $k$, $r$, and $\theta$ are the spring constant, equilibrium bond distance, and angle. The units of $m$, $R$, $k_r$, $k_\theta$, $r$ and $\theta$ are reported in amu, Å, kcal/mol·Å$^2$, kcal/mol·radian$^2$, Å and radian, respectively.

| Parameter | Value |
|---|---|
| $m_P$ | 78.9596 |
| $m_S$ | 117.0557 |
| $m_B$ | 134.0472 |
| $R_P$ | 1.89 |
| $R_S$ | 2.61 |
| $R_B$ | 2.52 |
| $k_{PP}$ | 23.0 |
| $k_{BS}$ | 10.0 |
| $k_{PS}$ | 23.0 |
| $k_{SP}$ | 64.0 |
| $k_{SS}$ | 23.0 |
| $r_0^{PP}$ | 3.3 |
| $r_0^{BS}$ | 4.8515 |
| $r_0^{PS}$ | 3.8157 |
| $r_0^{SP}$ | 4.601 |
| $r_0^{SS}$ | 2.7 |
| $k_{PPS}$ | 20.0 |
| $k_{BSP}$ | 5.0 |
| $k_{PSS}$ | 20.0 |
| $k_{BSS}$ | 5.0 |
| $k_{SSP_1}$ | 20.0 |
| $k_{SP_1P_1}$ | 20.0 |
| $\theta_0^{PPS}$ | 2.06 |
| $\theta_0^{BSP}$ | 1.7029 |
| $\theta_0^{PSS}$ | 1.444 |
| $\theta_0^{BSS}$ | 1.9259 |
| $\theta_0^{SSP_1}$ | 2.4 |
| $\theta_0^{SP_1P_1}$ | 1.4 |

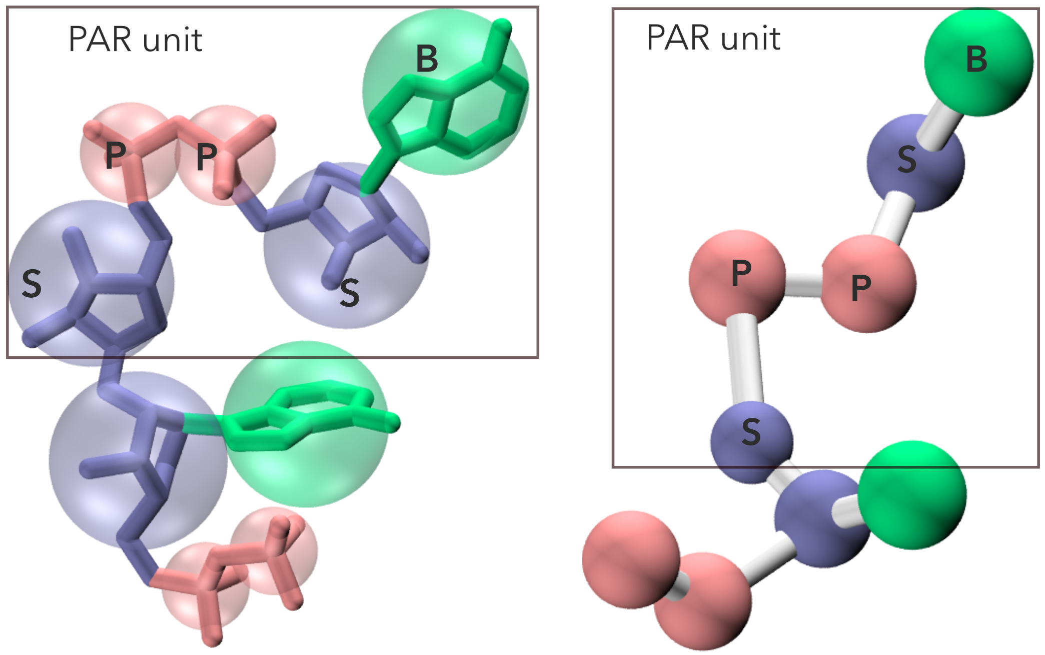


Figure S1: All-atom and coarse-grained representation of Poly(ADP-Ribose) (PAR), where one monomer of PAR is highlighted within a black box. Each monomer consists of five CG beads: two phosphate (P) groups, two sugar (S) groups, and one adenine base (B).

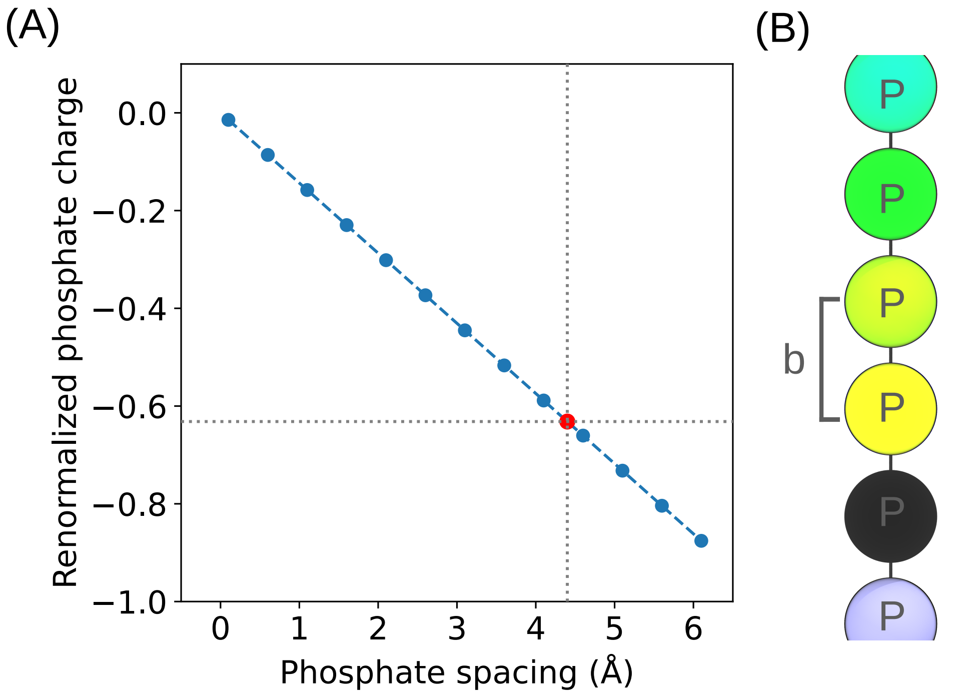


Figure S2: **(A)** $Q_P$ as a function of phosphate spacing ($b$) at 300 K. The renormalized value is $-Q_P = -\frac{b}{l_B}$, where $l_B$ is the Bjerrum length. The red dot indicate the value used in the simulations, $b = 4.4$ Å. **(B)** Illustration of $b$.

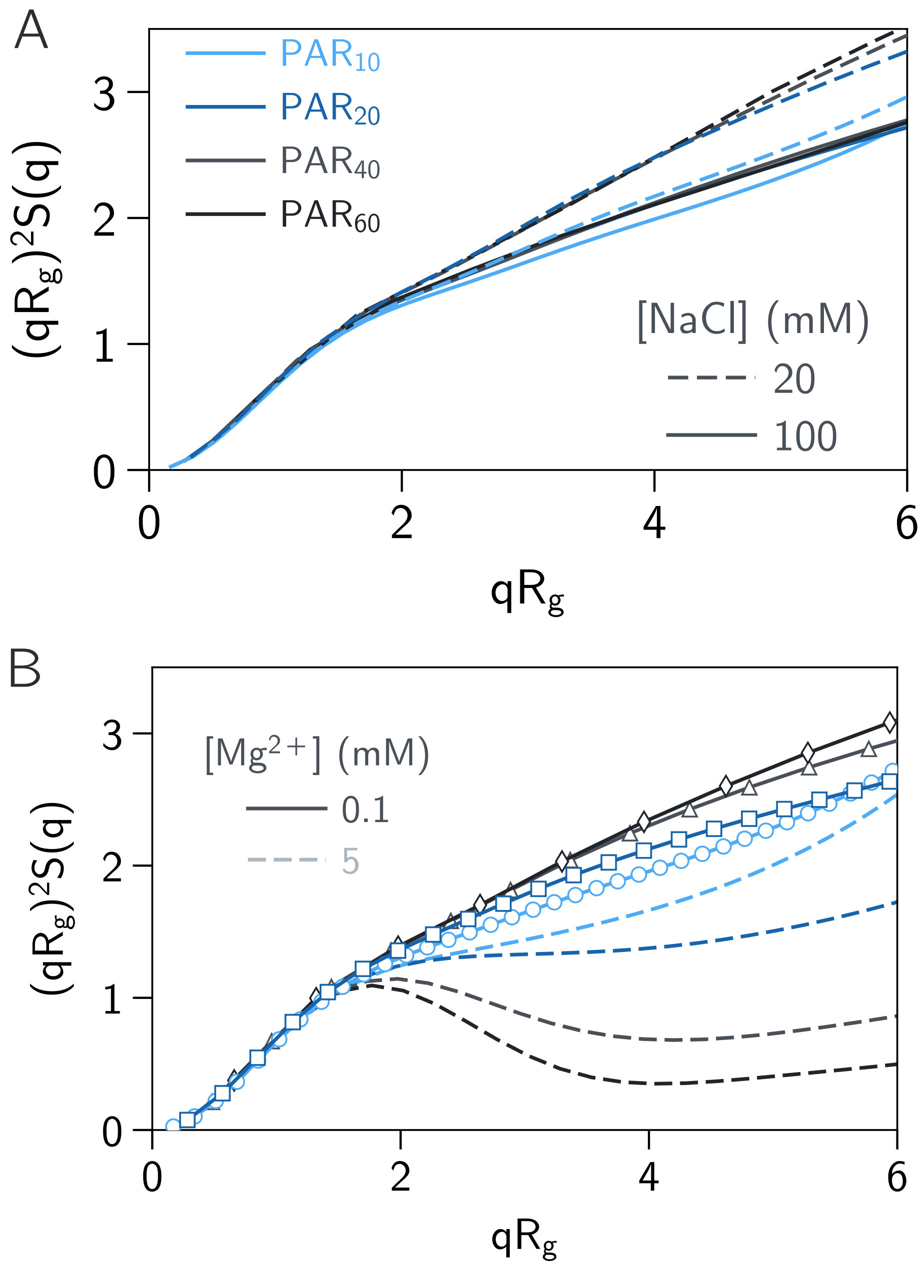


Figure S3: Dimensionless Kratky plots for different PAR lengths in (A) NaCl and (B) $Mg^{2+}$ ion solutions.

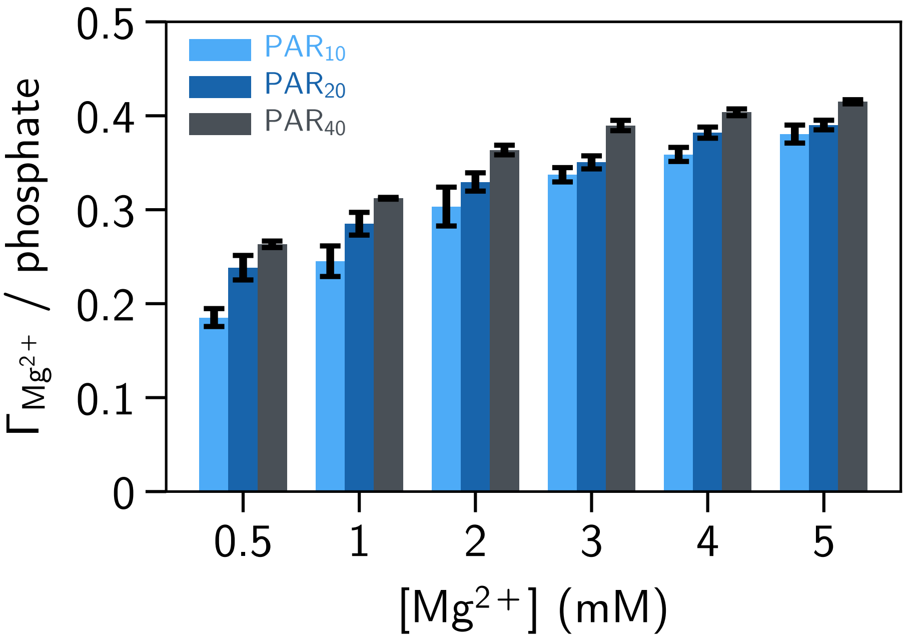


Figure S4: Excess $Mg^{2+}$ per phosphate with varying $Mg^{2+}$ concentrations for different PAR lengths.

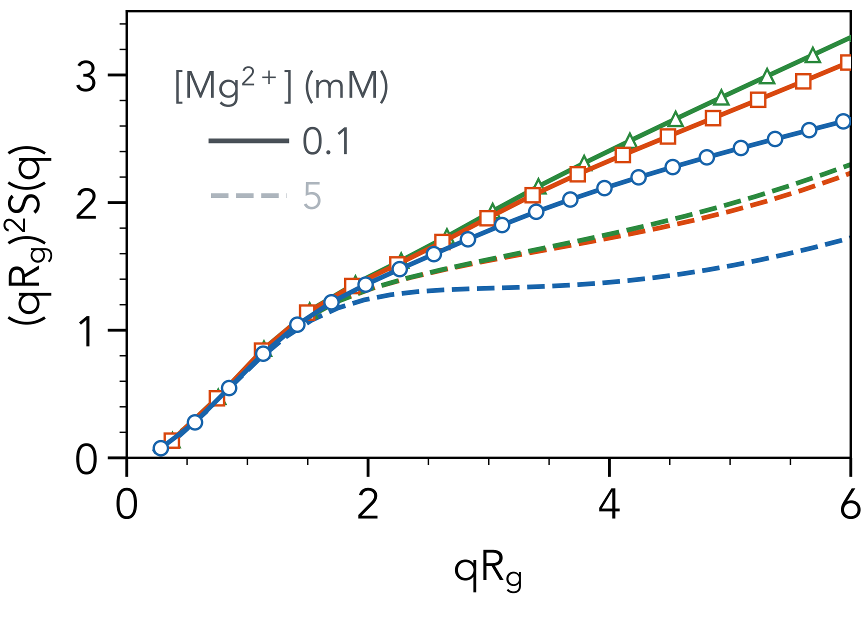


Figure S5: Dimensionless Kratky plots for $PAR_{20}$ (blue), $rA_{40}$ (orange) and $rU_{40}$ (green) at 0.1 mM (solid lines with marker) and 5 mM (dashed lines) $Mg^{2+}$.

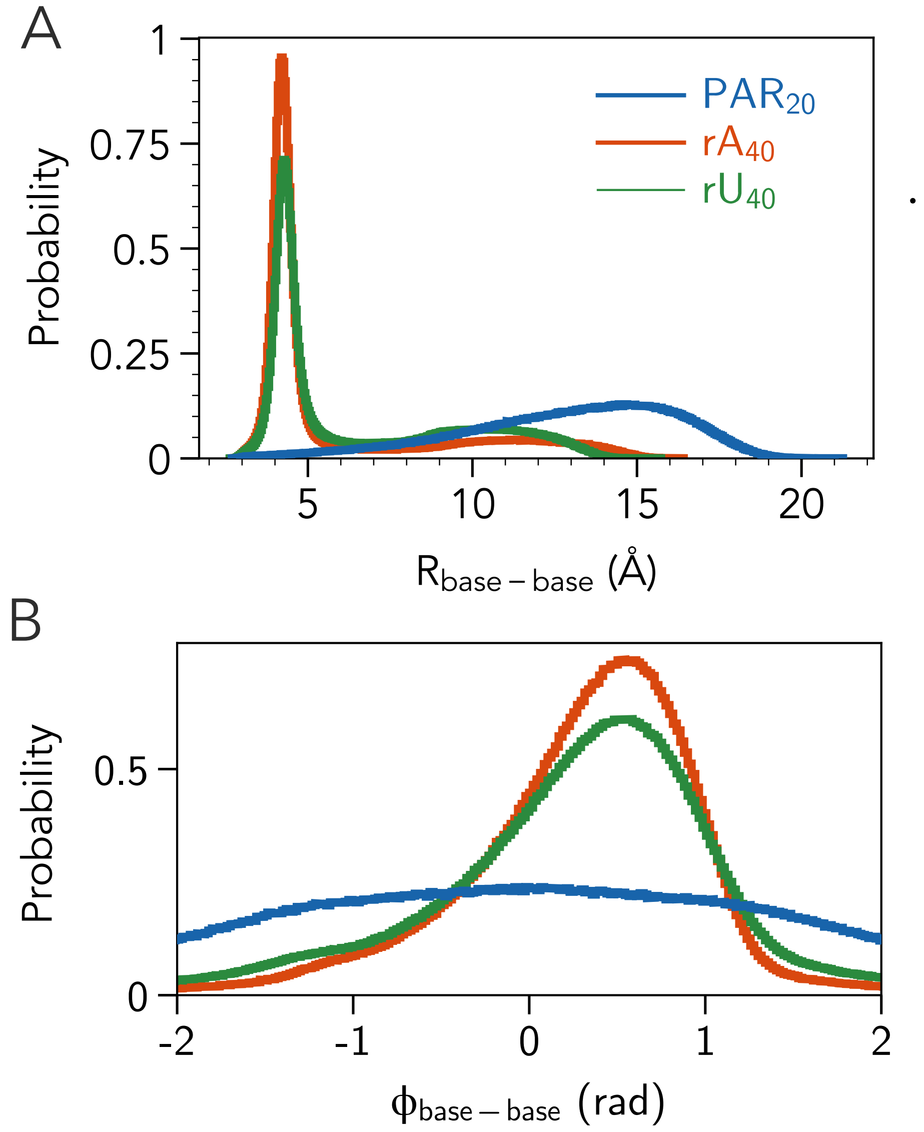


Figure S6: Consecutive base-base distance (top) and dihedral angle (bottom) in a mixture of 20 mM NaCl + 1 mM $Mg^{2+}$.

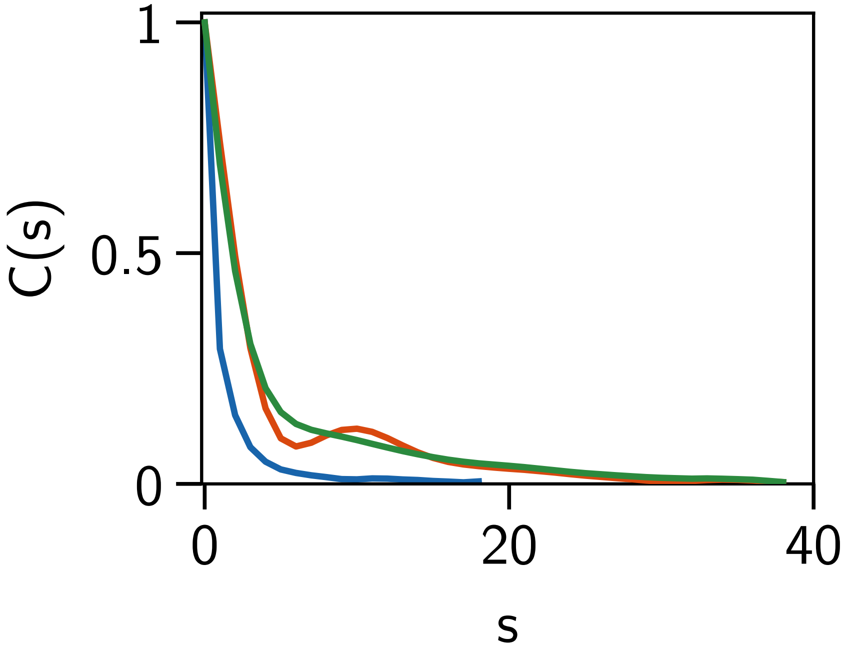


Figure S7: Bond autocorrelation for $PAR_{20}$ (blue), $rA_{40}$ (orange) and $rU_{40}$ (green) in a mixture of 20 mM NaCl + 1 mM $Mg^{2+}$.

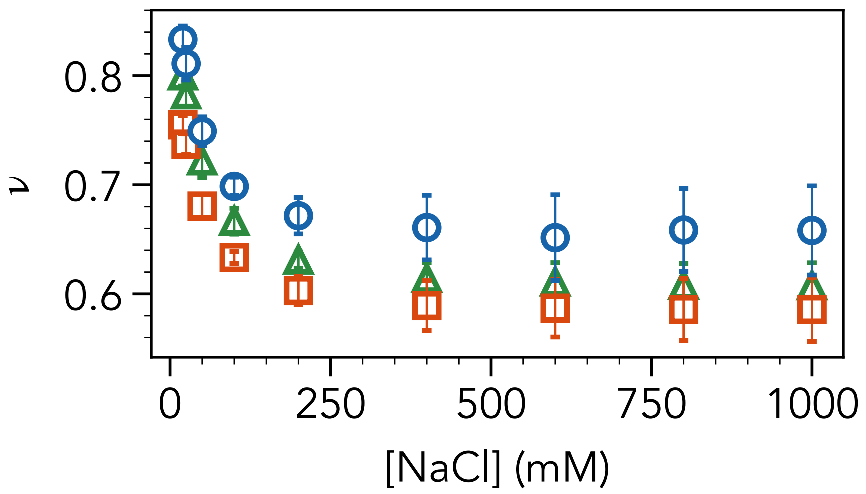


Figure S8: Apparent scaling exponent $\nu$ for $PAR_{20}$ (blue), $rA_{40}$ (orange) and $rU_{40}$ (green).

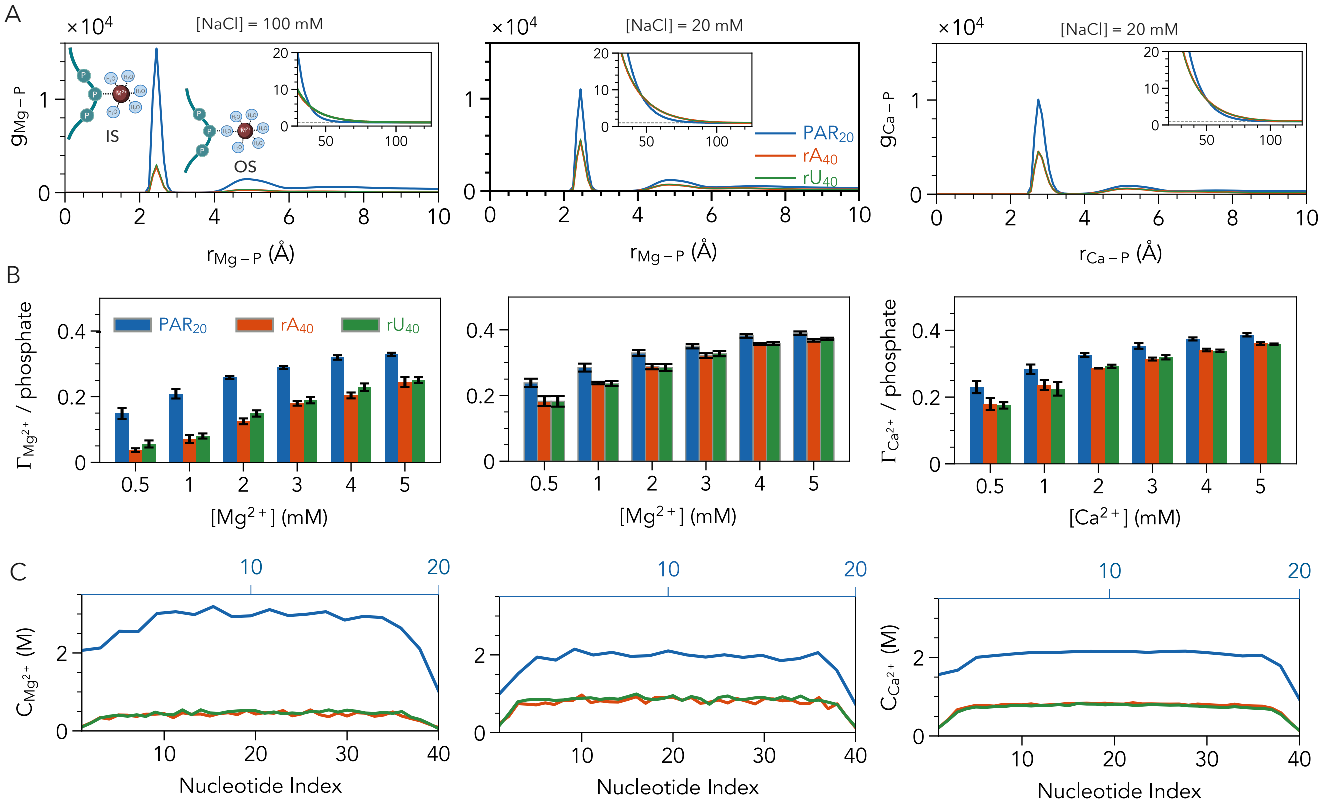


Figure S9: (A) Radial distribution function between $M^{2+}$ (M = Mg, Ca) ions and phosphate for $PAR_{20}$ (blue), $rA_{40}$ (orange), and $rU_{40}$ (green). The three panels correspond to different ionic conditions: (left) 100 mM NaCl + 1 mM $Mg^{2+}$, (middle) 20 mM NaCl + 1 mM $Mg^{2+}$, (right) 20 mM NaCl + 1 mM $Ca^{2+}$. The inset shows the decay of $g_{M-P}$ at large separation. (B) $\Gamma_{M^{2+}}$ for three molecules, with panels corresponding to the same ionic conditions in (A). (C) Local inner-sphere $M^{2+}$ ion concentration at each monomer unit for three molecules under the same ionic conditions in (A). The bottom and top labels on the x axis correspond to RNA and PAR unit indices, respectively.

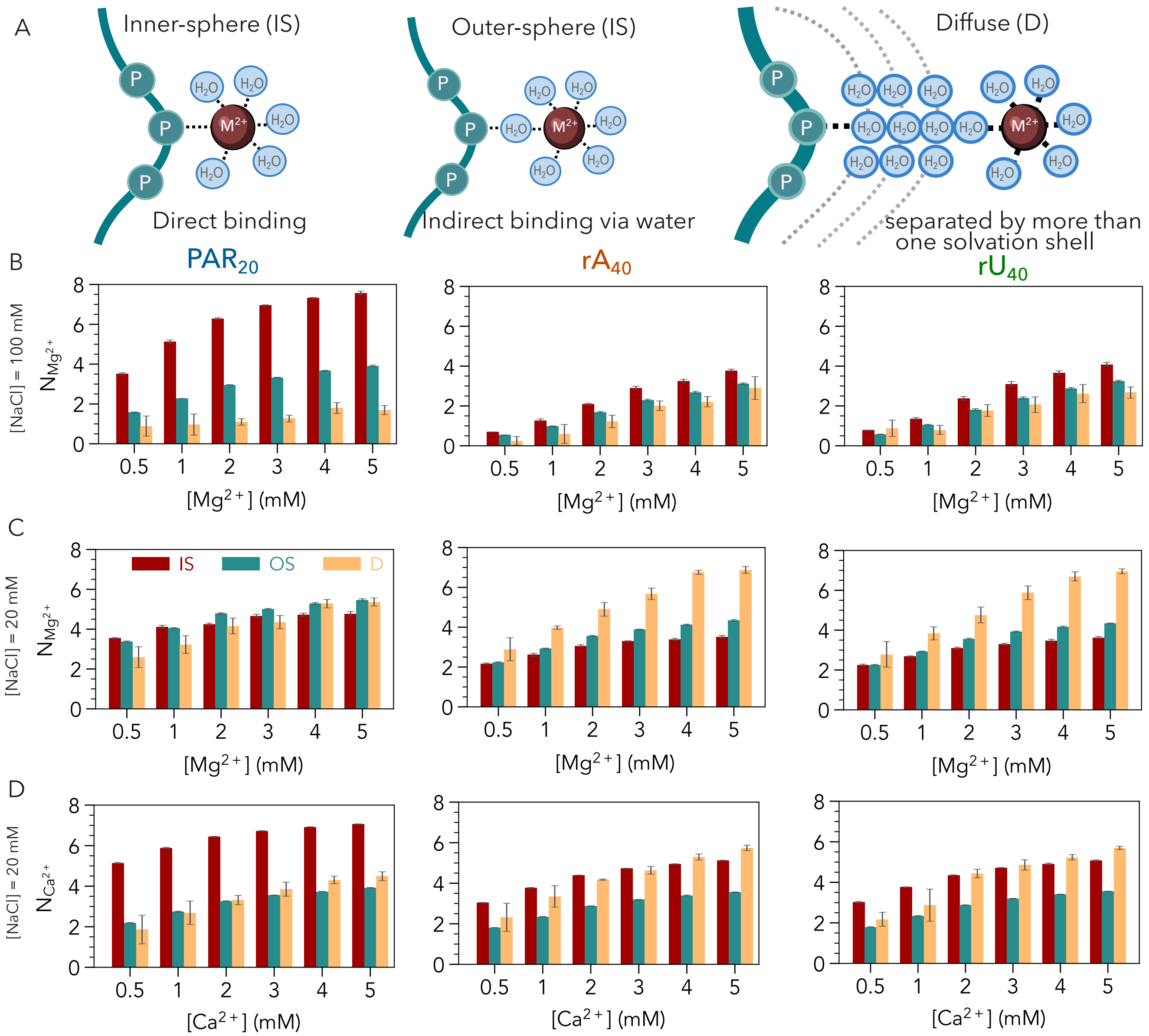


Figure S10: **Ion Atmosphere of PAR and RNA:** (A) Schematic diagram of ion binding modes: inner-shell (IS), outer-shell (OS), and diffusive (D). IS ions are in direct contact with phosphate whereas OS ions interaction is mediated by a single water molecule. Diffusive ions are separated from phosphate atoms by more than one solvation shells. Number of IS, OS, and diffuse ions, $N_{M^{2+}}$ (M = Mg, Ca) for (left) $PAR_{20}$, (middle) $rA_{40}$, and (right) $rU_{40}$ in a mixture of (B) 100 mM NaCl and varying $Mg^{2+}$, (C) 20 mM NaCl and varying $Mg^{2+}$ and (D) 20 mM NaCl and varying $Ca^{2+}$.

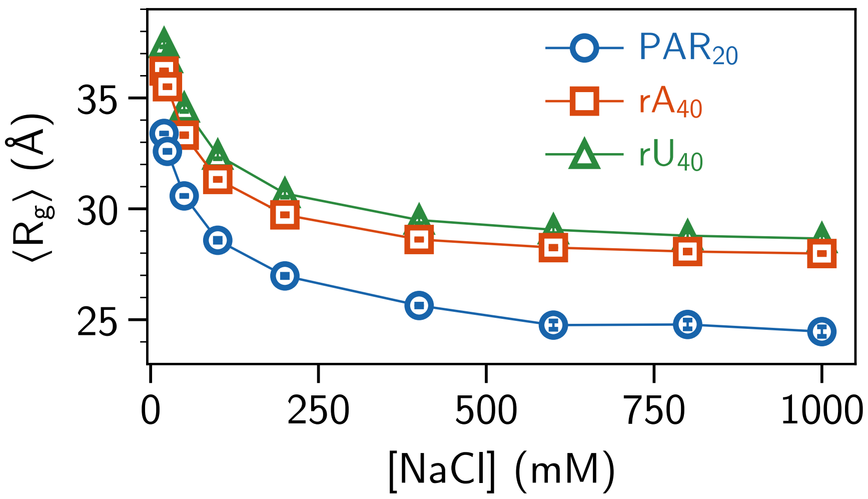


Figure S11: $R_g$ of $PAR_{20}$ (blue), $rA_{40}$ (orange) and $rU_{40}$ (green) in monovalent salt.

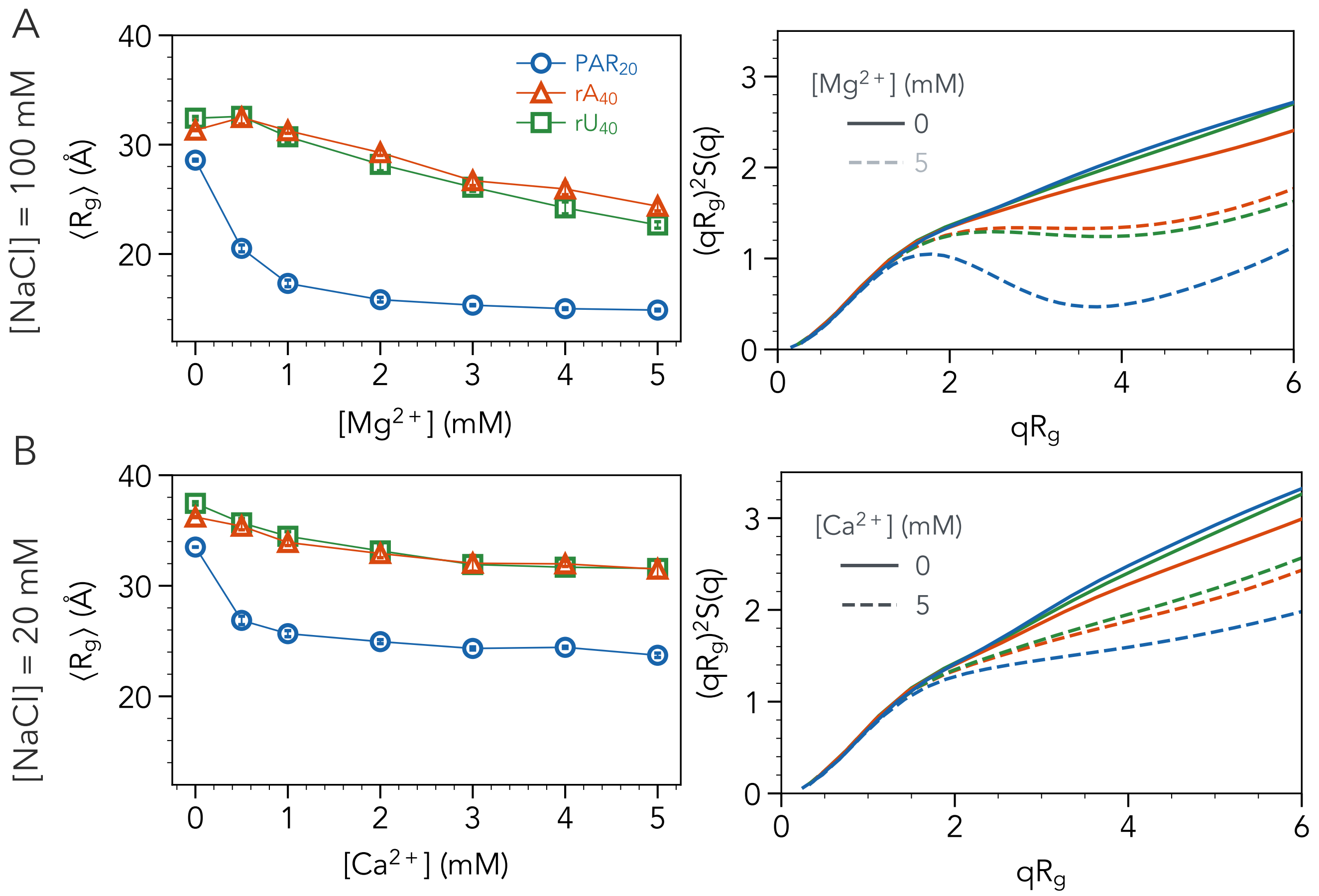


Figure S12: (A) (Left) $R_g$ of $PAR_{20}$ (blue circles), $rA_{40}$ (orange squares) and $rU_{40}$ (green triangles) in the presence of $Mg^{2+}$ (with 100 mM NaCl background). (Right) Dimensionless Kratky plots for three molecules without (solid lines) and with 5 mM $Mg^{2+}$ (dashed lines) (100 mM NaCl background). (B) Same as A, but for $Ca^{2+}$ in the presence of 20 mM NaCl.

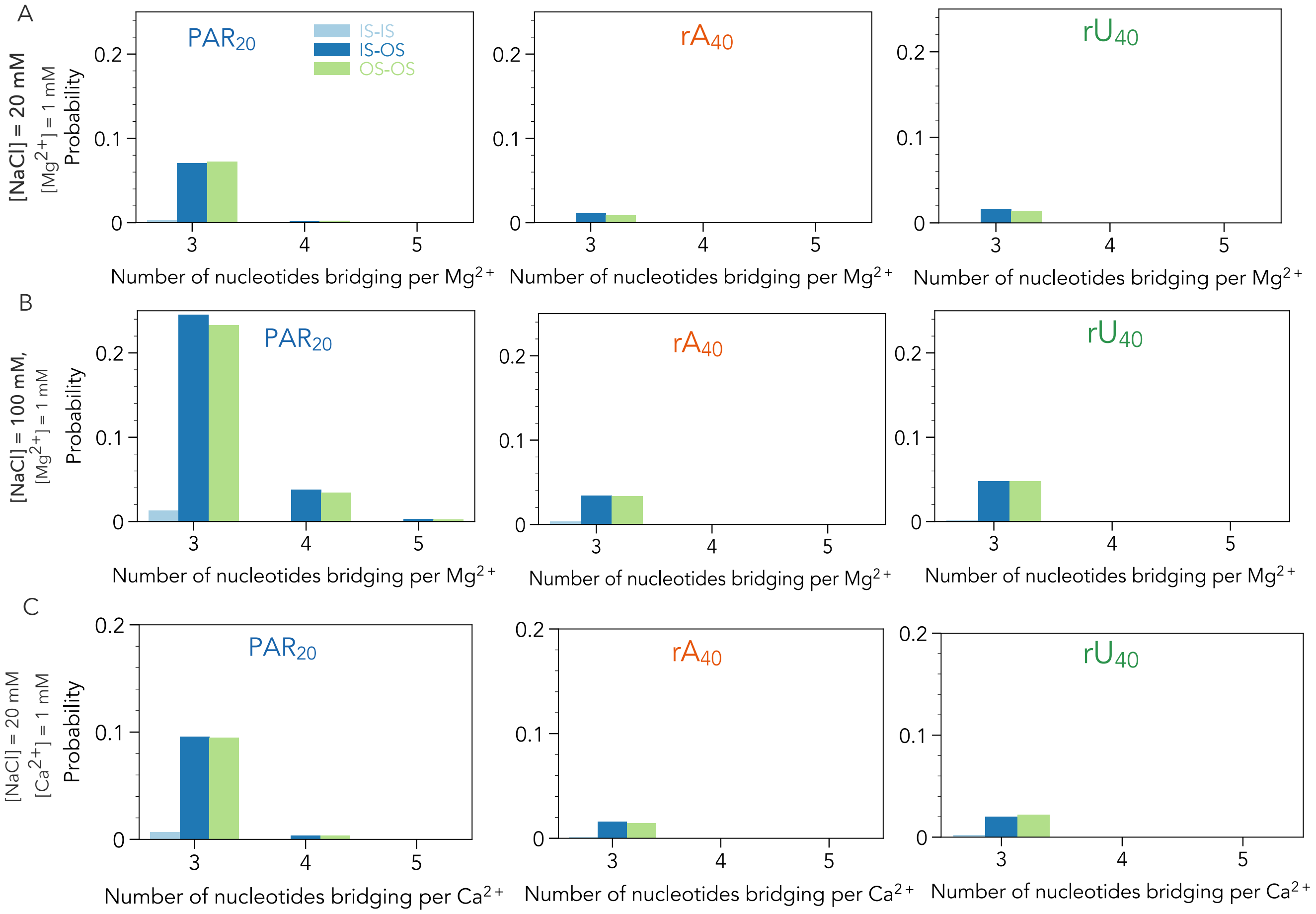


Figure S13: Fraction of bridging $n > 2$ phosphate groups simultaneously via IS-IS, IS-OS and OS-OS modes at (A) 20 mM NaCl + 1 mM $Mg^{2+}$, (B) 100 mM NaCl + 1 mM $Mg^{2+}$, and (C) 20 mM NaCl + 1 mM $Ca^{2+}$ for $PAR_{20}$, $rA_{40}$ and $rU_{40}$.

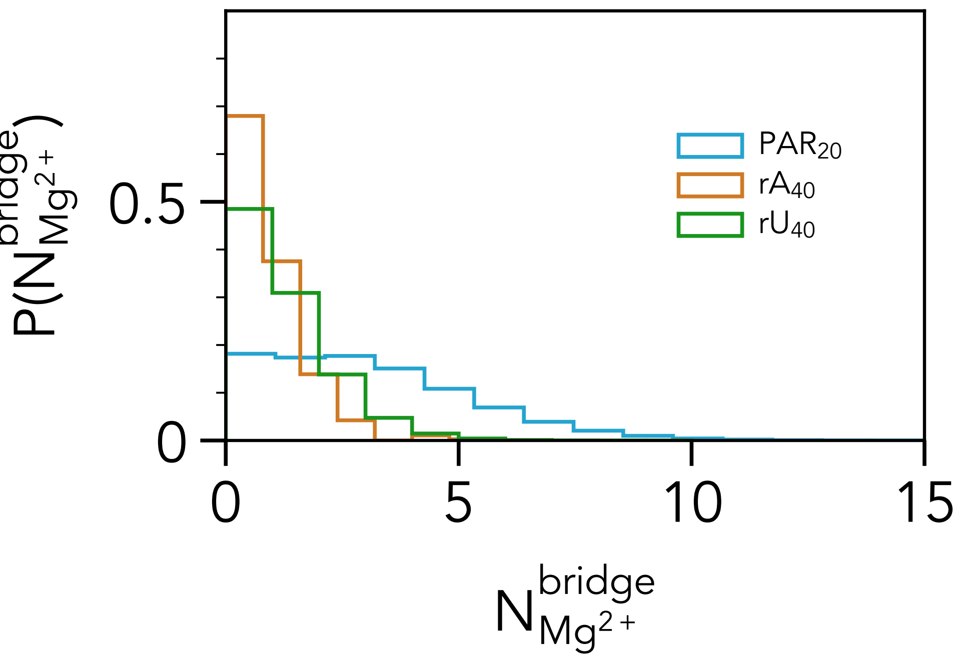


Figure S14: Distribution of number of bridging ions, $N_{Mg^{2+}}^{bridge}$ for $PAR_{20}$, $rA_{40}$ and $rU_{40}$ at 20 mM NaCl + 1 mM $Mg^{2+}$.

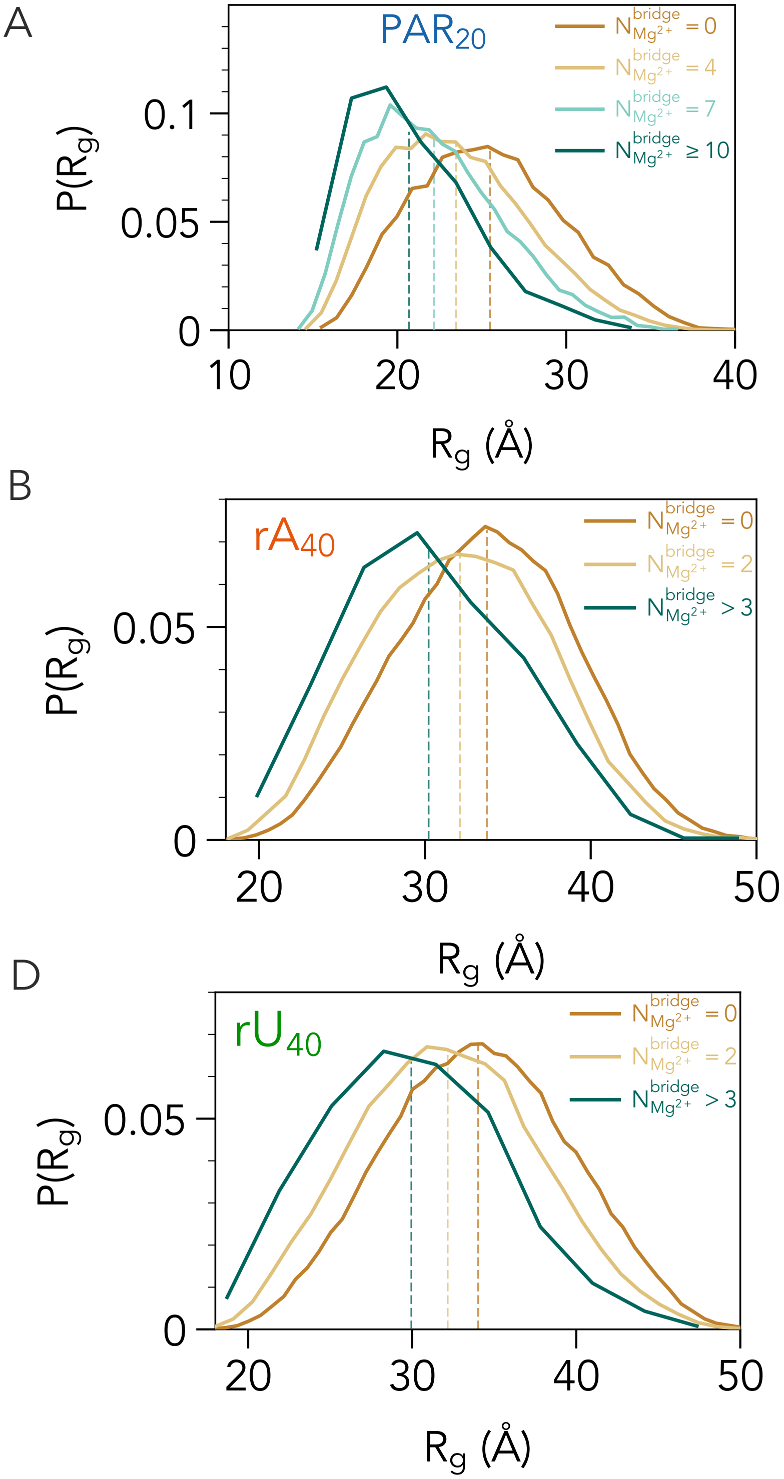


Figure S15: $R_g$ distribution for the conformations with $N^{bridge}_{Mg^{2+}}$ (= 0, 4, 7 and $\geq$ 7) for (A) $PAR_{20}$, (= 0, 2, > 3) for (B) $rA_{40}$ and (= 0, 2, > 3) for (C) $rU_{40}$ at 20 mM NaCl + 1 mM $Mg^{2+}$. The mean values are denoted by color-coded vertical lines.

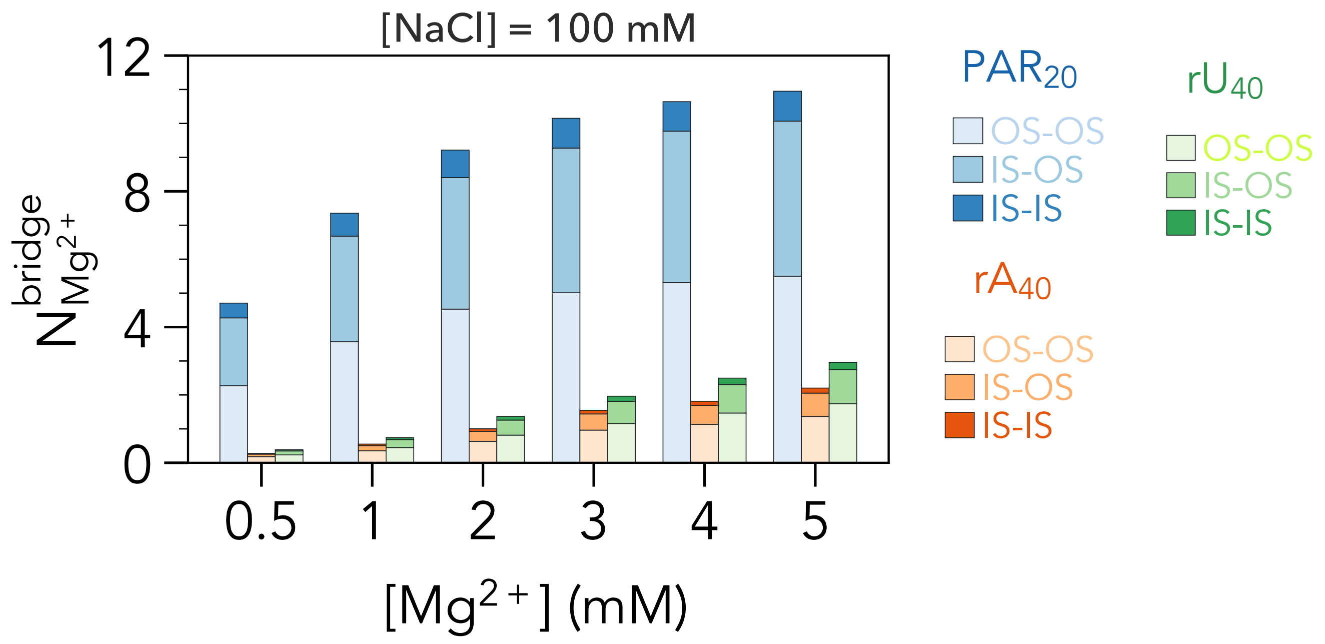


Figure S16: Number of bridging divalent ions via IS-IS, IS-OS, and OS-OS modes in a mixture of 100 mM NaCl + varying $Mg^{2+}$ for $PAR_{20}$ (blue), $rA_{40}$ (orange), and $rU_{40}$ (green).

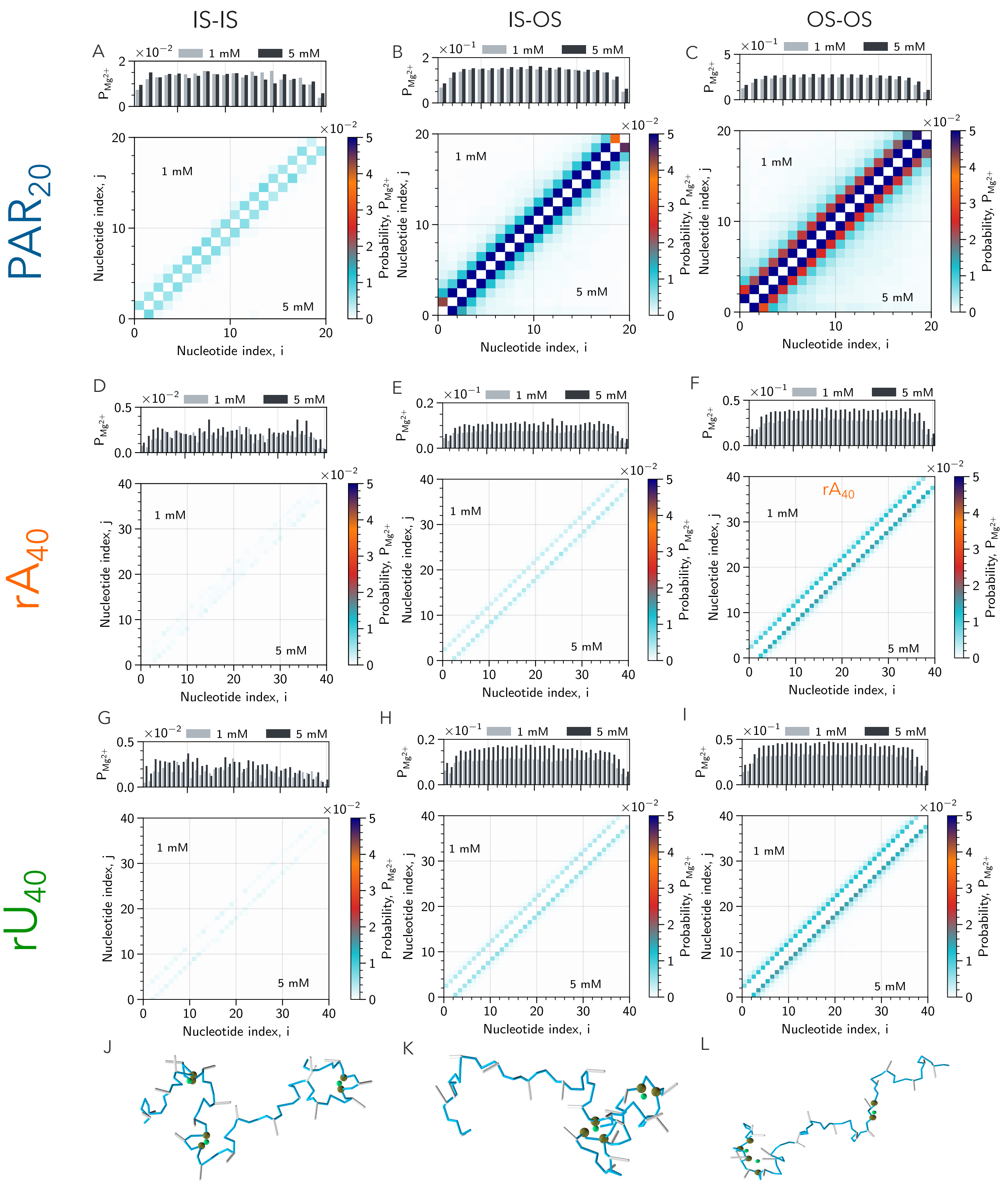


Figure S17: Probability of bridging between nucleotides by $Mg^{2+}$ ions in a mixture of 20 mM NaCl + 1 mM (upper panel) or 5 mM (lower panel) $Mg^{2+}$ for (A) IS-IS, (B) IS-OS, and (C) OS-OS bridging modes in $PAR_{20}$. The color bar represents the bridging probability, $P_{Mg^{2+}}$. The probability of bridging per nucleotide at 1 mM and 5 mM $Mg^{2+}$ is shown at the top. Similar data for $rA_{40}$ in (D), (E), and (F), and for $rU_{40}$ in (G), (H), and (I). Representative $PAR_{20}$ conformations with (J) IS-IS, (K) IS-OS, and (L) OS-OS bridging modes are shown in the bottom.

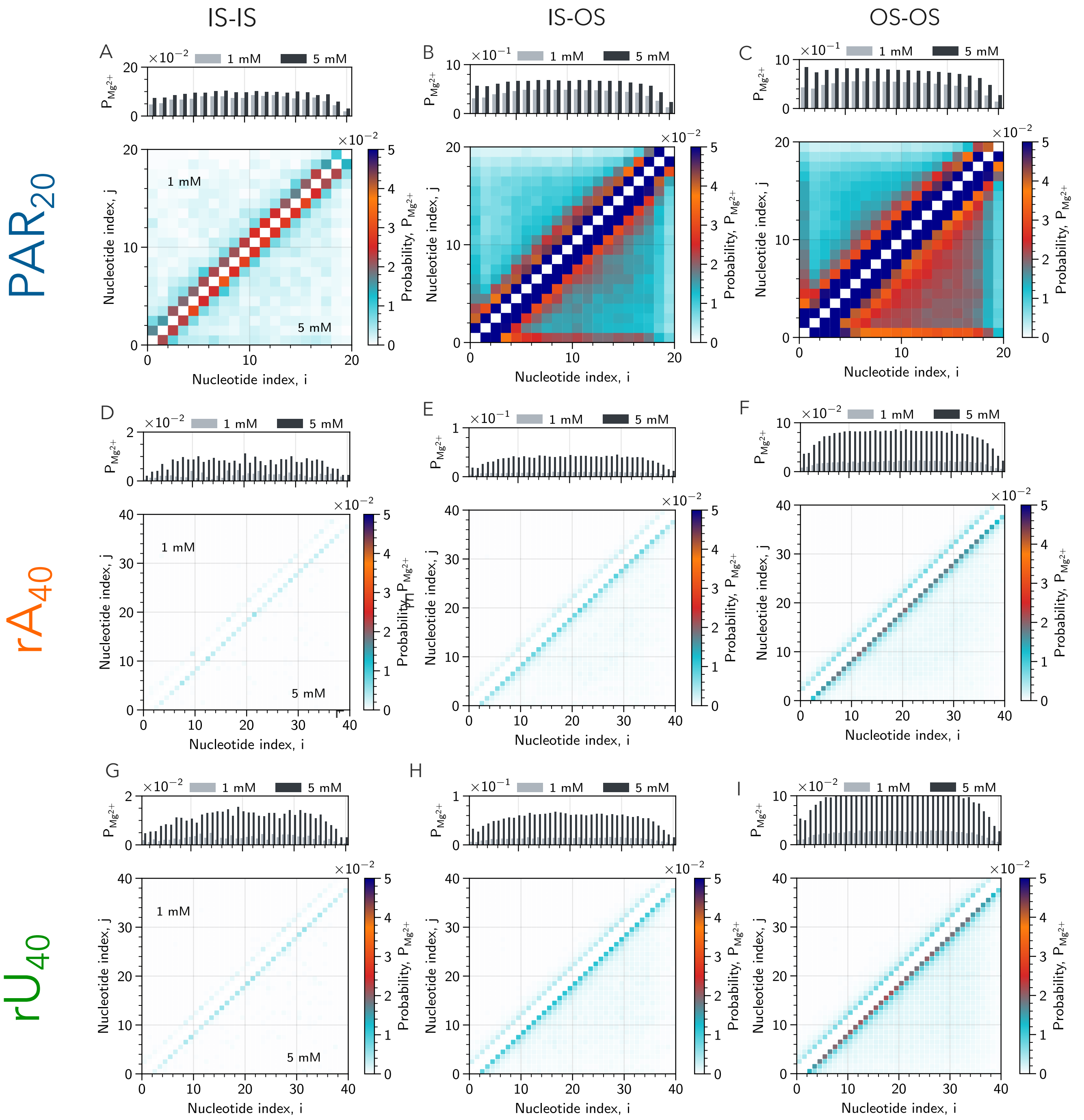


Figure S18: Same as Fig. S17, but at higher 100 mM NaCl.

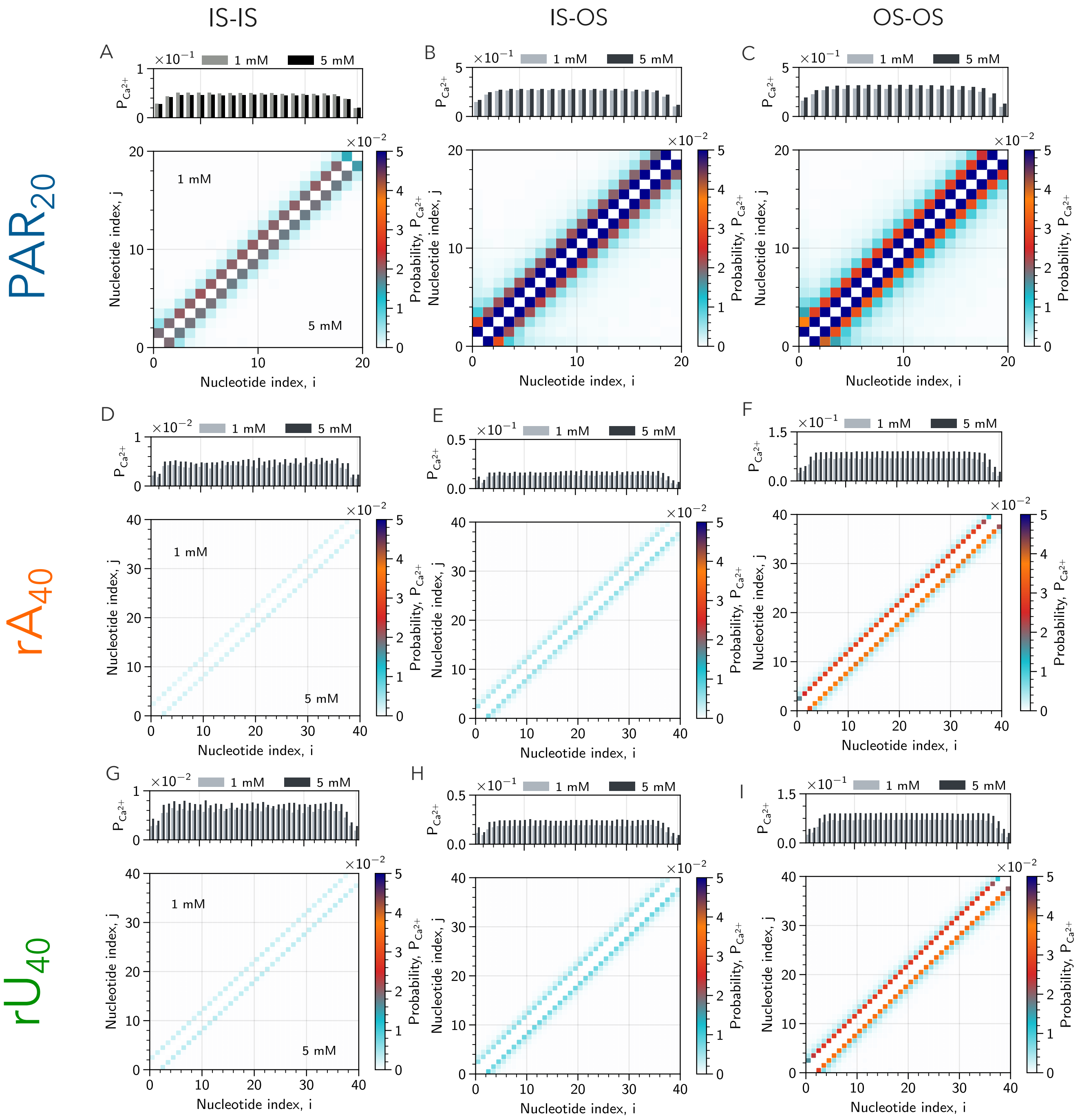


Figure S19: Same as Fig. S17, but for $Ca^{2+}$.

(9) Eastman, P.; Swails, J.; Chodera, J. D.; McGibbon, R. T.; Zhao, Y.; Beauchamp, K. A.; Wang, L.-P.; Simmonett, A. C.; Harrigan, M. P.; Stern, C. D. et al. OpenMM 7: Rapid

development of high performance algorithms for molecular dynamics. *PLoS Comput. Biol.* **2017**, *13*, e1005659.
